# A systematic survey of regional multitaxon biodiversity: evaluating strategies and coverage

**DOI:** 10.1101/158030

**Authors:** Ane Kirstine Brunbjerg, Hans Henrik Bruun, Lars Brøndum, Aimée T. Classen, Lars Dalby, Kåre Fog, Tobias G. Frøslev, Irina Goldberg, Anders Johannes Hansen, Morten D.D. Hansen, Toke T. Høye, Anders A. Illum, Thomas Læssøe, Gregory S. Newman, Lars Skipper, Ulrik Søchting, Rasmus Ejrnæs

## Abstract

**Background:** In light of the biodiversity crisis and our limited ability to explain variation in biodiversity, tools to quantify spatial and temporal variation in biodiversity and its underlying drivers are critically needed. Inspired by the recently published ecospace framework, we developed and tested a sampling design for environmental and biotic mapping. We selected 130 study sites (40 × 40 m) across Denmark using stratified random sampling along the major environmental gradients underlying biotic variation. Using standardized methods, we collected site species data on vascular plants, bryophytes, macrofungi, lichens, gastropods and arthropods. To evaluate sampling efficiency, we calculated regional coverage (relative to the known species number per taxonomic group), and site scale coverage (i.e., sample completeness per taxonomic group at each site). To extend taxonomic coverage to organisms that are difficult to sample by classical inventories (e.g., nematodes and non-fruiting fungi), we collected soil for metabarcoding. Finally, to assess site conditions, we mapped abiotic conditions, biotic resources and habitat continuity.

**Results:** Despite the 130 study sites only covering a minute fraction (0.0005 %) of the total Danish terrestrial area, we found 1774 species of macrofungi (54 % of the Danish fungal species pool), 663 vascular plant species (42 %), 254 bryophyte species (41 %) and 200 lichen species (19 %). For arthropods, we observed 330 spider species (58 %), 123 carabid beetle species (37 %) and 99 hoverfly species (33 %). Correlations among species richness for taxonomic groups were predominantly positive. Overall, sample coverage was remarkably high across taxonomic groups and sufficient to capture substantial spatial variation in biodiversity across Denmark. This inventory is nationally unprecedented in detail and resulted in the discovery of 143 species with no previous record for Denmark. Comparison between plant OTUs detected in soil DNA and observed plant species confirmed the usefulness of carefully curated environmental DNA-data. Species richness did not correlate well among taxa suggesting differential and complex biotic responses to environmental variation.

**Conclusions:** We successfully and adequately sampled a wide range of diverse taxa along key environmental gradients across Denmark using an approach that includes multi-taxon biodiversity assessment and ecospace mapping. Our approach is applicable to assessments of biodiversity in other regions and biomes where species are structured along environmental gradient.

## Background

The vast number of species on Earth have yet to be described, challenging our understanding of biodiversity [1]. For a deeper understanding of what determines the distribution of species across the planet, comprehensive data on species occurrence and environmental conditions are required. While some progress has been made in understanding the distribution of biodiversity at coarse spatial resolution, our knowledge of biodiversity at high spatial resolution is deficient [2]. In this study, we consider biodiversity as the richness and spatial turnover of taxonomic units, whether species or operational taxonomic units (OTUs) derived by eDNA (environmental DNA) metabarcoding. While progress has been made in the interpretation and prediction of richness and turnover of vascular plants and vertebrates, various types of bias, e.g. temporal, spatial, and taxonomic bias [3], have constrained similar advances for less well-known, but diverse groups such as fungi and insects [1]. As a result, conservation management is typically based on biodiversity data from a non-random subset of taxa [4].

Recent developments in molecular techniques – in particular the extraction and sequencing of eDNA – hold the promise of more time-efficient sampling and identification of species [5, 6]. Further, eDNA enables the exploration of communities and organisms not easily recorded by traditional biodiversity assessment, such as soil-dwelling nematodes [7]. In fact, PCR-based methods combined with DNA sequencing have already provided valuable insight into the taxonomic diversity within complex environmental samples, such as soil [8–10] and water [e.g. 11, 12]. Due to the ongoing rapid development in DNA sequencing technologies, with the emergence of next generation sequencing (NGS) techniques − generating billions of DNA sequences [13] − an environmental sample could now be analyzed to a molecular depth that gives an almost exhaustive picture of the species composition at the site of collection. Despite this potential, rigorous assessments with complete taxonomic coverage from eDNA samples are still missing [6, 12, 14]. To assess the suitability and potential of eDNA data in complementing − or even replacing − traditional field survey data, tests on comprehensive data sets are needed.

Undertaking an ambitious biodiversity field study across a wide geographical space comes with major logistical and methodological challenges. It is not clear what environmental gradients structure biodiversity across the tree of life and for most taxa standardized field protocols to sample species occurrences are non-existent. The recently developed ecospace framework suggests that biodiversity varies in relation to its position along environmental gradients (position), the availability of biotic resources, such as organic matter and structures e.g. trees for epiphytes (expansion), and spatio-temporal extent of biotopes (continuity) [15]. Environmental conditions and local processes can be a template shaping local biodiversity (e.g. through environmental filtering) [16, 17]. This template is highlighted by the ecospace position of sampled biotopes in abiotic environmental space. In addition to the physico-chemical conditions shaping abiotic gradients − particularly important to autotrophic organisms − the presence and abundance of specific biological resources, crucial to heterotrophic organisms, such as specialist herbivores, detritivores and saproxylic species are likely important and thus should be considered [18]. The quantification of biotic resources and structures, e.g. dead wood, dung and carcasses, is not often included in community studies, despite the limited knowledge in the area [15, 19] which speaks for further studies. Spatial and temporal processes at regional extent, such as extinction, speciation and migration, shape species pools and thereby set the limits to local richness and species composition [16, 17, 20]. In order to improve our understanding of biodiversity patterns, local and regional factors should be considered concurrently [17, 21].

In this study, we used the ecospace framework as guideline to develop a comprehensive sampling design for large-scale mapping of variation in biodiversity and environmental variation across Denmark. Data collection and analysis was carried out as part of a research project (called *Biowide*).

The fundamental and radical claim of ecospace is that a low-dimensional environmental hyperspace can be used to predict and forecast variation in multi-taxon species richness. Testing this claim demands a dataset covering the variation of the terrestrial environment in a region of significant spatial coverage, a representative sample of the major taxonomic groups contributing to α-diversity and a mapping of the most important environmental factors defining the conditions for the terrestrial biota.

The project aimed to cover all of the major environmental gradients, including variation in soil moisture, soil fertility and succession, as well as habitats under cultivation. Within this environmental space spanned by 130 40 × 40 m sites, we performed a systematic and comprehensive sampling of the environment and biodiversity. We combined traditional species observation and identification with modern methods of biodiversity mapping in the form of massive parallel sequencing of eDNA extracted from soil samples.

In this paper we present the inventory and evaluate whether we achieved the comprehensive and representative data collection needed to test the claimed generality of the ecospace framework.

## Methods

### Study area and site selection

We aimed to characterize biodiversity across the country of Denmark (Fig. 1a) – a lowland area of 42,934 km^2^ and an elevational range of 0-200 meters above sea level. While there are some limestone and chalk outcrops, there is no exposed bedrock in the investigated area. Soil texture ranges from coarse sands to heavy clay and organic soils of various origins [22]. Land use is dominated by arable land (61 %), most of which is in annual rotation, while forests are mostly plantations established during the 19^th^ and 20^th^ centuries. Scrubs cover approximately 17 %, natural and semi-natural terrestrial habitats some 10 %, and freshwater lakes and streams 2 %. The remaining 10 % is made up of urban areas and infrastructure [23, 24].

**Figure 1:**
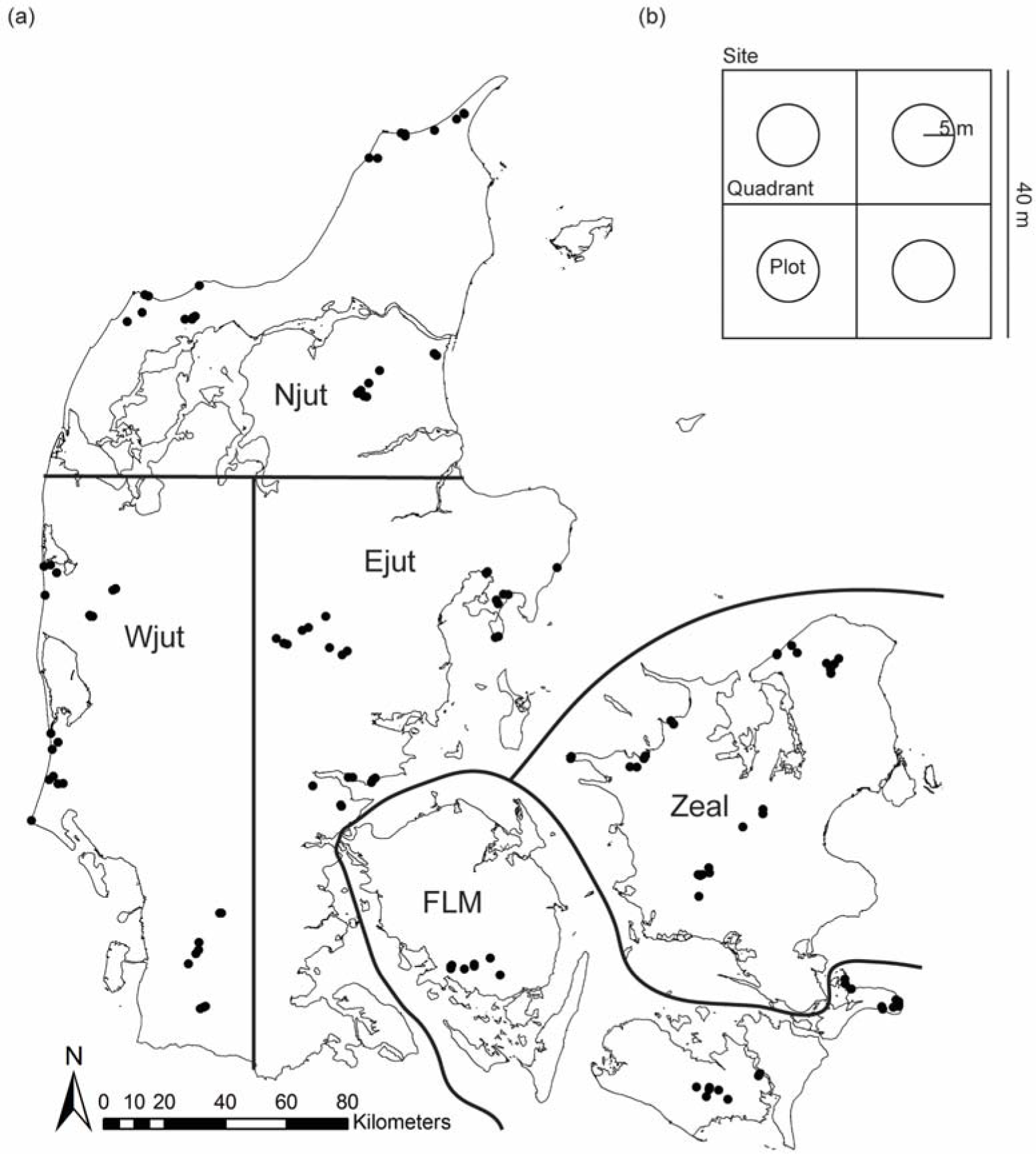
a) Map of Denmark showing the location of the 130 sites grouped into 15 clusters within five regions (Njut: Northern Jutland, Wjut: Western Jutland, Ejut: Eastern Jutland, FLM: Funen, Lolland, Møn, Zeal: Zealand). b) Site layout with four 20 × 20 m quadrants each containing a 5 m radius circle (plot).

When selecting sites, we considered major environmental gradients, the potential size of the sampling units (sites), as well as practicalities of sampling across the large geographical space within the same season. The sites were 40 × 40 m which was a compromise between within-site homogeneity and the representativeness of a particular habitat type. We stratified site selection according to the identified major environmental gradients, including the intensity of human land use. We measured 30 sites that were cultivated habitats and 100 sites that were natural and semi-natural habitats. This balance between natural and cultivated habitat was chosen, because we expected cultivated habitats to have shorter environmental gradients. The cultivated subset represented major land-use categories and the natural subset was stratified across natural gradients in soil fertility, soil moisture, and successional stage from sparsely vegetated to closed canopy forest, (Appendix A). We deliberately excluded linear features, such as hedgerows and road verges, urban areas with predominantly exotic plants as well as saline and aquatic habitats, but included temporarily inundated heath, dune depressions and wet mires.

The final set of 25 sampling classes consisted of six cultivated habitat types; three types of fields (rotational, leys, and oldfield) and three types of plantations (beech, oak, and spruce). 18 natural classes consisted of all factorial combinations of natural soil fertility (fertile or infertile), moisture (dry, moist, or wet), and successional stage (low vegetation with bare soil, closed herb/scrub, or forest) (Appendix A). Finally, we included a class of perceived areas of high species richness [25] in Denmark. These sites were selected subjectively by performing a public poll among active natural history volunteers in the Danish nature conservation and nature management societies. The 25 classes were replicated in each of five geographical regions within Denmark (Fig. 1a). The result was 130 sites with 18 natural, 6 cultivated, and two perceived areas of high species richness evenly distributed across each of five geographic regions of Denmark (Table 1). For logistical reasons, we did not place any sites on Bornholm although we acknowledge that this island is geologically different than the rest of Denmark.

**Table 1:** Stratification of sites in the survey. The sites are sub-divided into four categories (arable, plantations, perceived areas of high species richness (HighSpcRich), and natural). The natural sites were stratified across specific levels of succession (early, mid, and late), soil moisture (wet, moist, and dry) and soil fertility (rich and poor), while this was the case for the other classes of sites. The number of sites within each of the 25 classes is given.

| Category | Class | Successional stage | Moisture | Fertility | Number of sites |
| --- | --- | --- | --- | --- | --- |
| Arable | Rotational | - | - | - | 5 |
| Arable | Ley | - | - | - | 5 |
| Arable | Old field | - | - | - | 5 |
| Plantation | Beech | - | - | - | 5 |
| Plantation | Oak | - | - | - | 5 |
| Plantation | Spruce | - | - | - | 5 |
| HighSpcRich | HighSpcRich | - | - | - | 10 |
| Natural | Early/Dry/Rich | Early | Dry | Rich | 5 |
| Natural | Mid/Dry/Rich | Mid | Dry | Rich | 5 |
| Natural | Late/Dry/Rich | Late | Dry | Rich | 5 |
| Natural | Early/Moist/Rich | Early | Moist | Rich | 5 |
| Natural | Mid/Moist/Rich | Mid | Moist | Rich | 5 |
| Natural | Late/Moist/Rich | Late | Moist | Rich | 5 |
| Natural | Early/Wet/Rich | Early | Wet | Rich | 5 |
| Natural | Mid/Wet/Rich | Mid | Wet | Rich | 5 |
| Natural | Late/Wet/Rich | Late | Wet | Rich | 5 |
| Natural | Early/Dry/Poor | Early | Dry | Poor | 5 |
| Natural | Mid/Dry/Poor | Mid | Dry | Poor | 5 |
| Natural | Late/Dry/Poor | Late | Dry | Poor | 5 |
| Natural | Early/Moist/Poor | Early | Moist | Poor | 5 |
| Natural | Mid/Moist/Poor | Mid | Moist | Poor | 5 |
| Natural | Late/Moist/Poor | Late | Moist | Poor | 5 |
| Natural | Early/Wet/Poor | Early | Wet | Poor | 5 |
| Natural | Mid/Wet/Poor | Mid | Wet | Poor | 5 |
| Natural | Late/Wet/Poor | Late | Wet | Poor | 5 |

For the 18 natural habitat classes, site selection through stratified random sampling was guided by a large nation-wide dataset of vegetation plots in semi-natural habitats distributed across the entire country (n = 96,400 plots of 78.5 m^2^ each, www.naturdata.dk) from a national monitoring and mapping project [26] and in accordance with the EU Habitats Directive [27]. We used environmental conditions computed from plant indicator values to select candidate sites for each class. First, we calculated plot mean values for Ellenberg indicator values based on vascular plants species lists [28] and Grime CSR-strategy allocations of recorded plants [29], the latter were recoded to numeric values following Ejrnæs & Bruun [30]. We excluded saline and artificially fertilized habitats by excluding plots with Ellenberg S > 1 or Ellenberg N > 6. We then defined stratification categories as: fertile (Ellenberg N 3.5-6.0), infertile (Ellenberg N < 3.5), dry (Ellenberg F < 5.5), moist (Ellenberg F 5.5-7.0), wet (Ellenberg F > 7.0), early succession (Grime R > 4 and Ellenberg L > 7 or > 10 % of annual plants), late succession (mapped as forest), mid succession (remaining sites).

To reduce transport time and costs, all 26 sites within each region were grouped into three geographic clusters (Fig. 1a). The nested sampling design allowed us to take spatially structured species distributions into account [31]. The procedure for site selection involved the following steps:

1. Designation of three geographic clusters within each region with the aim to cover all natural classes while a) keeping the cluster area below 200 km^2^ and b) ensuring high between-cluster dispersion in order to represent the geographic range of the region. In practice, perceived areas of high species richness were chosen first, then clusters were placed with reference to the highest ranking areas of high species richness and in areas with a wide range of classes represented in the national vegetation plot data [32].
2. Representing the remaining 24 classes in each region by selecting 8-9 potential sites in each cluster. Sites representing natural classes were selected from vegetation plot data. Cultivated classes were assumed omnipresent and used as buffers in the process of completing the non-trivial task of finding all classes within each of three cluster areas of < 200 km^2^ in each region.
3. Negotiating with land owners and, in case of disagreement, replacing the preferred site with an alternative site from the same class.

After each of the 130 sites were selected using available data, we established each 40 × 40 m site in a subjectively selected homogenous area that accounted for topography and vegetation structure. Each site was divided into four 20 × 20 m quadrants, and from the center of each quadrant a 5 m radius circle (called a plot) was used as a sub-unit for data collection to supplement the data collected at site level (40 × 40 m) (Fig. 1b).

### Collection of biodiversity data

For each of the 130 sites, we aimed at making an unbiased and representative assessment of multi-taxon species richness. Data on vascular plants, bryophytes, lichens, macrofungi, arthropods and gastropods were collected using standard field inventory methods (Appendix B). For vascular plants, bryophytes and gastropods, we collected exhaustive species lists. For the remaining taxonomic groups that are more demanding to find, catch, and identify, we aimed at collecting a reproducible and unbiased sample through a standardized level of effort (typically one hour). Multiple substrates (soil, herbaceous debris, wood, stone surfaces and bark of trees up to 2 m) were carefully searched for lichens and macrofungi at each site. For fungi, we visited each site twice during the main fruiting season in 2014 – in August and early November – and once during the main fruiting season in 2015 – between late August and early October. Specimens that were not possible to identify with certainty in the field were sampled and, when possible, identified in the laboratory. For arthropod sampling, a standard set of pitfall traps (including meat-baited and dung-baited traps), yellow Möricke pan traps and Malaise traps were operated during a fixed period of the year. In addition, we used active search and collection methods, including sweep netting and beating as well as expert searches for plant gallers, miners and gastropods. Finally, we heat-extracted collembolas and oribatid mites from soil cores. Due to the limited size of the sites relative to the mobility of mammals, birds, reptiles and amphibians, data on these groups were not recorded. Records of arthropods were entered in https://www.naturbasen.dk. Records of fungi were entered in https://svampe.databasen.org/. All species occurrence data and environmental data has been made available at the project home page http://bios.au.dk/om-instituttet/organisation/biodiversitet/projekter/biowide/. Species data will be made available for GBIF (www.gbif.org) through the above-mentioned web portals. Specimens are stored at the Natural History Museum Aarhus (fungal specimens at the fungarium at the Natural History Museum of Denmark). For further details on the methods used for collection of biodiversity data see Appendix B.

### Collection of eDNA data

We used soil samples collected from all 130 sites for the eDNA inventory. At each site, we sampled 81 soil cores in a 9 × 9 grid covering the entire 40 m × 40 m plot and pooled the collected samples after removal of coarse litter. We homogenized the soil by mixing with a mixing paddle mounted on a drilling machine. A subsample of soil was sampled from the homogenized sample and DNA was extracted for marker gene amplification and sequencing [14]. We chose the MiSeq platform by Illumina for DNA sequencing. MiSeq is adapted to amplicon sequencing [33]. For further details on methods for eDNA data generation and considerations on eDNA species richness and community composition measures see Appendix B.

Data from the fungal eDNA community matrix was mapped to the Darwin Core data standard (http://rs.tdwg.org/dwc/) and wrapped in a DwC archive for publication to the Global Biodiversity Information Facility. The ‘dataGeneralizations’ field was used to indicate the identity of OTUs towards the UNITE species hypothesis concept [34], Sampling sites were included as WKT polygons in the ‘footprintWKT’ field and sampling site names were included in the ‘eventID’ field. The representative sequences (OTUs) were included using the GGBN amplification extension. The dataset is available from gbif.org (https://doi.org/10.15468/nesbvx).

### Site environmental data

We have followed the suggestion in Brunbjerg et al. [15] to describe the fundamental requirements for biodiversity in terms of the ecospace (position, expansion and spatio-temporal continuity of the biotope).

#### Position

To assess the environmental variation across the 130 sites, we measured a core set of site factors that described the abiotic conditions at each site. Environmental recordings and estimates included soil pH, total soil carbon (C, g/m^2^), total soil nitrogen (N, g/m^2^) and total soil phosphorus (P, g/m^2^), soil moisture (% volumetric water content), leaf CNP (%), soil surface temperature (°C) and humidity (vapour pressure deficit), air temperature (°C), light intensity (Lux), and boulder density. For further details on methods used to collect abiotic data see Appendix B.

#### Expansion

We collected measurements that represent the expansion or biotic resources which some species consume and the organic and inorganic structures which some species use as habitat. Although many invertebrates are associated with other animals, for practical reasons, we restricted our quantification of biotic resources to the variation in live and dead plant tissue, including dung. We measured litter mass (g/m^2^), plant species richness, vegetation height (of herb layer, cm), cover of bare soil (%), bryophyte cover (%) and lichen cover (%), dead wood volume (m^3^/site), dominant herbs, the abundance of woody species, the number of woody plant individuals, flower density (basic distance abundance estimate, [35]), density of dung (basic distance abundance estimate), number of carcasses, fine woody debris density (basic distance abundance estimate), ant nest density (basic distance abundance estimate), and water puddle density (basic distance abundance estimate). For further details on methods used to collect expansion data see Appendix B.

#### Mapping of temporal and spatial continuity

For each site, we inspected a temporal sequence of aerial photos (from 1945 to 2014) and historical maps (1842-1945) starting with the most recent photo taken. We defined temporal continuity as the number of years since the most recent major documented land use change. The year in which a change was identified was recorded as a ‘break in continuity’. To estimate spatial continuity, we used ArcGIS to construct four buffers for each site (500 m, 1000 m, 2000 m, 5000 m). Within each buffer we estimated the amount of habitat similar to the site focal habitat by visual inspection of aerial photos with overlays representing nation-wide mapping of semi-natural habitat. For further details on methods for collection of continuity data see Appendix B.

### Analyses

To illustrate the coverage of the three main gradients (moisture, fertility, and successional stage) spanned by the 130 sites, Ellenberg mean site values (mean of mean Ellenberg values for the four 5m radius quadrats within each site) for soil moisture (Ellenberg F), soil nutrients (Ellenberg N) and light conditions (Ellenberg L) were plotted relative to Ellenberg F, N and L values for a reference data set of 5 m radius vegetation quadrats (47202 from agricultural, semi-natural and natural open vegetation and 12014 from forests (www.naturdata.dk) [26]. Mean Ellenberg values were only calculated for quadrats with more than five species and 95 percentile convex hull polygons where drawn for the reference data set as well as the Biowide data set.

We assessed the coverage for each taxonomic group across sites as well as within each site for spiders, harvestmen, and insect orders represented by at minimum of 75 species, for which we had abundance data by comparing the number of species found to the estimated species richness of the sample using rarefaction in the iNEXT R-package [36]. Coverage regarding habitat types was assessed by constructing species accumulation curves for arable sites, plantations and natural sites. To visualize the habitat type related differences in expansion (biotic resources) we created a radar chart illustrating flower density, dead wood volume, plant richness, litter mass and dung density for natural habitats (early, mid, late succession), plantations and arable sites.

To further evaluate the turnover component of biodiversity and how well we covered the environmental gradient for our inventory, we related community composition to the measured environmental variables (abiotic and biotic) based on a Nonmetric Multidimensional Scaling (NMDS) analyses in R v. 3.2.3 [37] using the vegan R-package [38] and the plant species × site matrix as well as the macrofungi species × site matrix. Abiotic and biotic variables were correlated with ordination axes to facilitate interpretation. In order to ensure that geographical variation in species composition and diversity was adequately assessed, we calculated, for each geographical region, the relative proportion of major species groups, and the component of beta diversity nestedness and turnover. The latter was done using the betapart R-package [39]. To illustrate and substantiate the adequacy of the eDNA sampling design and subsequent laboratory protocols, we correlated basic biodiversity measures of community composition (NMDS axes) and richness for plant eDNA (ITS2 marker region) with the same measures for our observed plant data (see Appendix B for detailed methods). To illustrate the cross correlation among the main taxonomic groups spearman rank correlations for vascular plants, mosses, lichens, macrofungi, gastropods, gallers/miners and arthropods at order level were calculated.

## Results

The 130 sites were distributed in 15 clusters nested within five regions across Denmark (Fig. 1a). The measured variables differed according to the initial stratification of sites based on simple indicators (Table 1, Fig. 2a, b, ranges of measured variables in Appendix C). Managed sites (plantations and agricultural fields) revealed little variation in soil moisture (Fig. 2b). The perceived areas of high species richness spanned the full variation of natural sites regarding fertility, moisture and successional stage (Fig. 2b).

**Figure 2:**
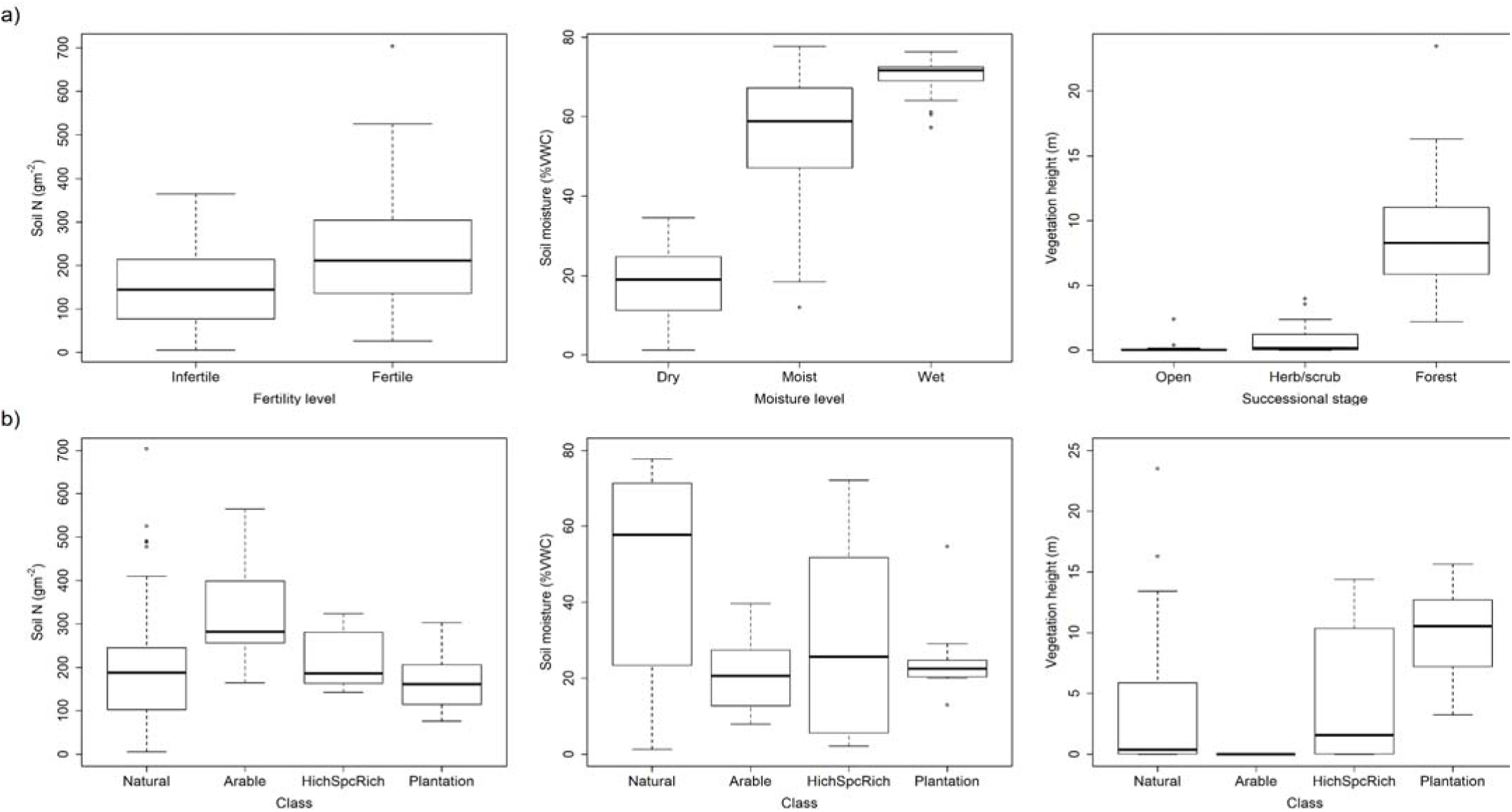
Validation of the stratification scheme used in site selection. Boxplots of measured values of nutrient levels (soil N g/m^2^), moisture levels (trimmed site mean % Volumetric Water Content (VWC)), and vegetation height (mean LIDAR canopy height (m)) for the a) 90 natural sites of different fertility levels (infertile, fertile), moisture levels (dry, moist, wet), and successional stages (early (open), mid (herb/scrub), late (forest)) and b) the 90 natural sites, 15 plantations, 15 fields and 10 perceived areas of high species richness (HighSpcRich).

The selected 130 sites covered the main gradients reflected by a huge reference dataset from a national monitoring program (Fig. 3) as judged from a vegetation-based calibration of site conditions regarding moisture, fertility and succession (light intensity). Biowide data seemed to increase the upper range of the fertility gradient, which can be explained by the inclusion in Biowide of rotational fields that were not included in reference data (Fig. 2b, 3).

**Figure 3:**
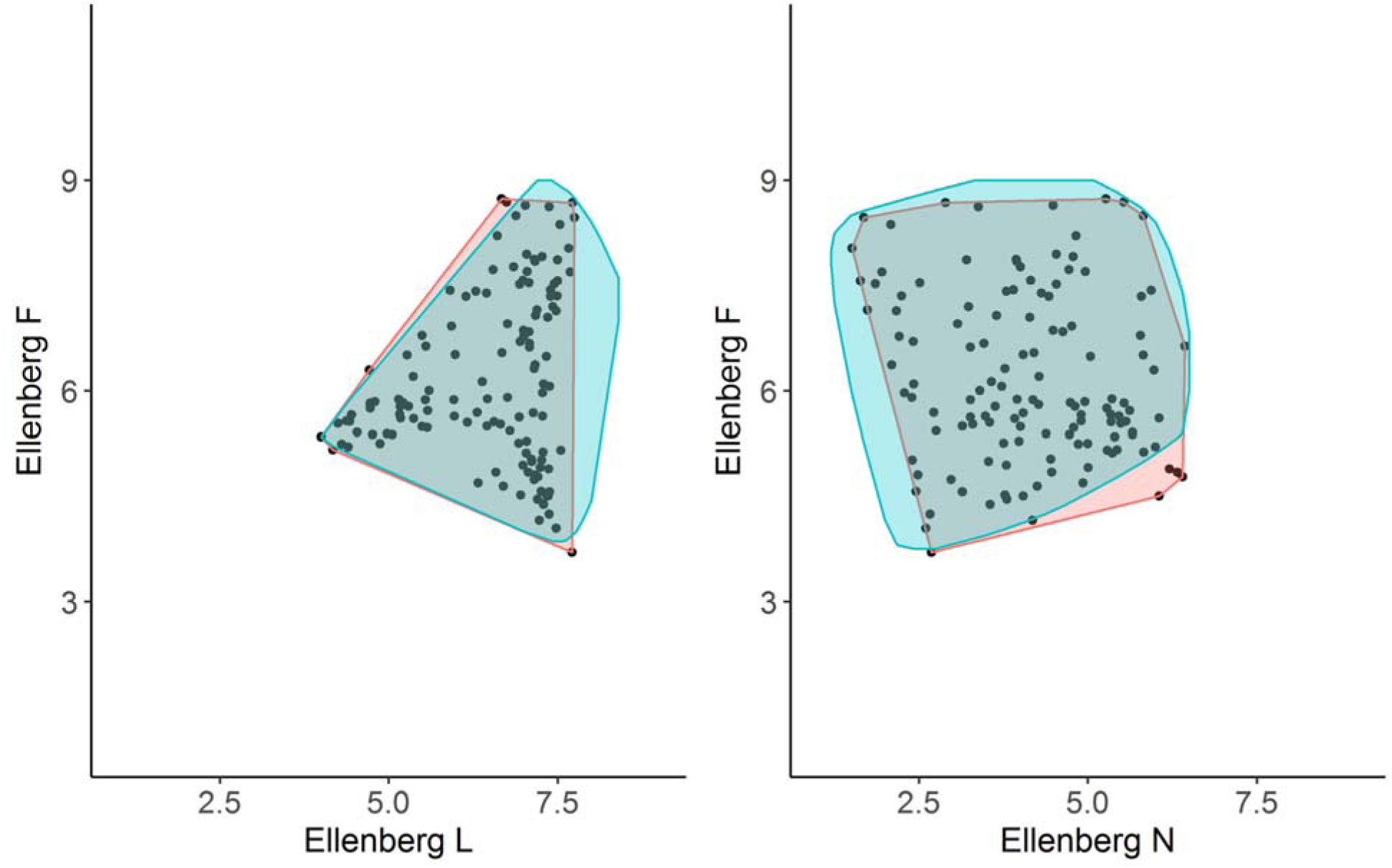
95 percentile convex hull plots of Ellenberg F, L and N values from a reference data set (www.naturdata.dk) of open and forest habitat types (blue, n= 59 227) as well as the data set used in this study, Biowide (red, n=130). Black dots represent Ellenberg values of the 130 Biowide sites.

The environmental expansion of ecospace, which was measured as the amount and differentiation of organic carbon sources, varied among habitat types with high litter mass in tree plantations and late successional habitats, high plant species richness in early and mid-successional habitats, high dung density in open habitats (early successional and fields) and high amounts of dead wood in late successional habitats (Fig. 4). Spatial and temporal continuity varied for the 130 sites with less spatial continuity at larger buffer sizes (Appendix D). The number of species found per site differed with taxonomic group with the highest number for arthropods and macrofungi and lowest for gastropods and lichens (Appendix E). There was no clear difference in relative richness, nestedness and turnover of taxonomic groups across geographic region.

**Figure 4:**
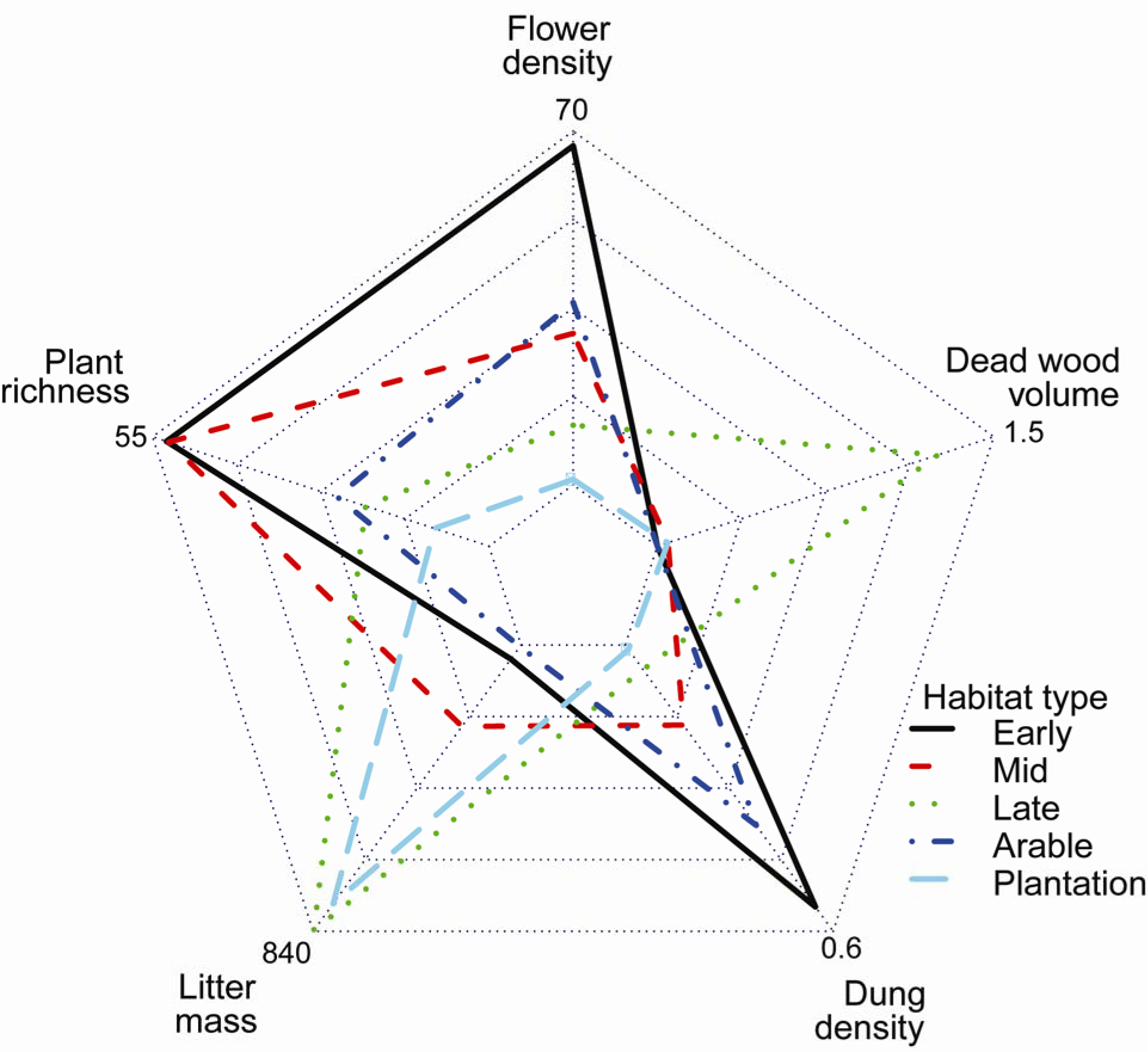
Habitat mean values for various carbon resources in the 130 40 × 40 m sites. Volume of dead wood (m^3^/ha), density of dung (cow, sheep, deer, horse, hare) (number/m^2^), summed flower density in April, June and August (number/m^2^), litter mass (g/m^2^) and plant species richness per site are depicted for natural habitat types (early, mid and late successional stage), arable sites and plantations.

We collected 1774 species of macrofungi (corresponding to 54 % of the number of macrofungi recorded in Denmark), 200 lichens (19 %), 663 vascular plants (42 %) and 254 bryophytes (41 %) during the study period. We collected 75 species of gastropods (75 %), 330 spiders (58 %), 99 hoverflies (33 %), 123 carabid beetles (37 %) and 203 gallers and miners species (21 %). For all groups except macrofungi, the number of species found was higher in natural (n = 90) than in cultivated (n = 40) sites, but across taxonomic groups, plantations and agricultural fields harbored species not found in other habitat types – plantations were particularly important in harboring unique species of macrofungi (Table 2, Appendix F). The taxonomic sample coverage calculated by rarefaction within the 130 sites was high overall (range: 0.86-0.99), but highest for gastropods and spiders and lowest for gallers and miners (Table 2). Species accumulation curves for three habitat categories (arable, plantation and natural) saturated at approximately the same high level (Appendix G).

**Table 2:**
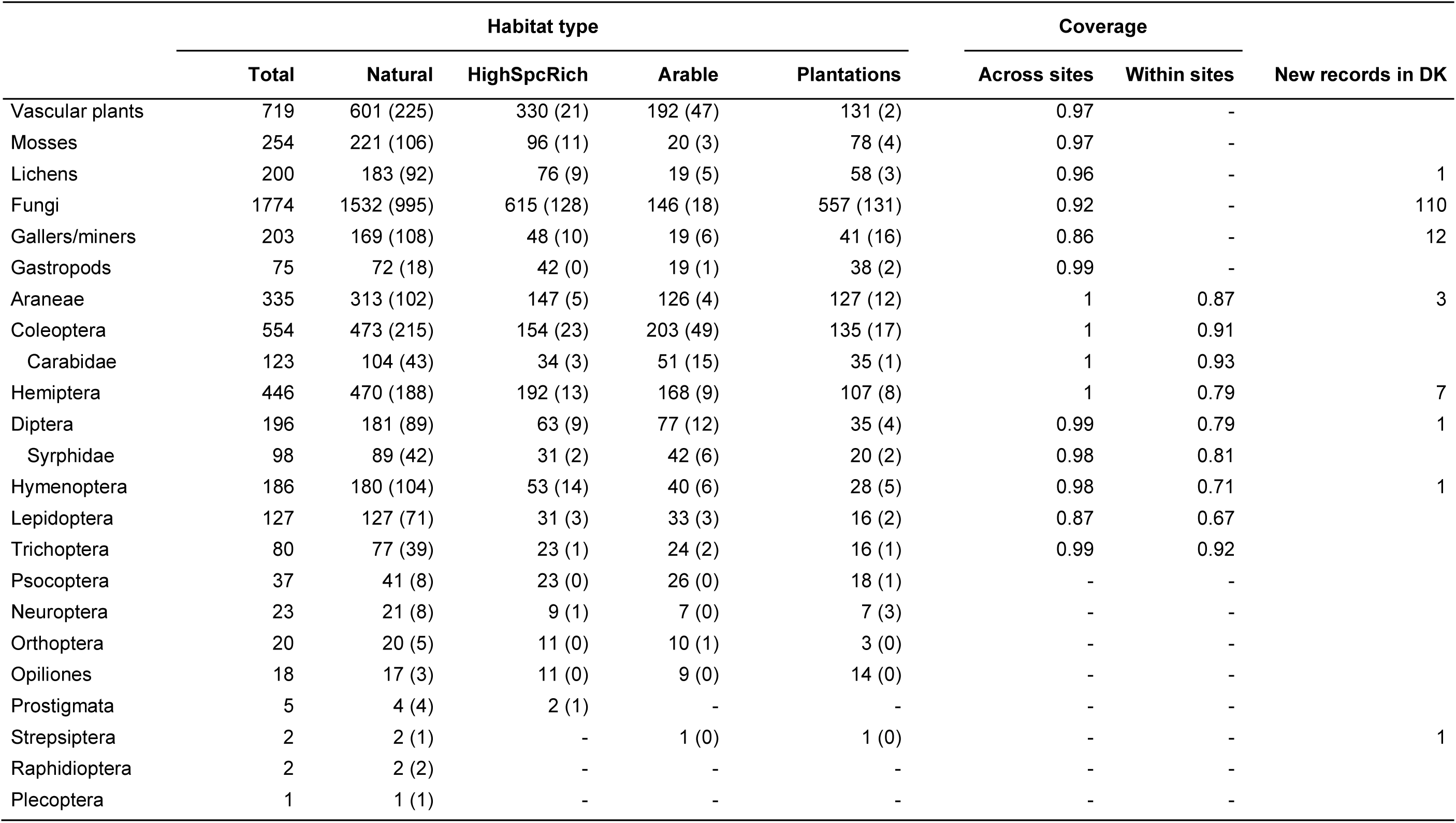
Species richness and sample coverage across habitats per taxonomic group. Number of species per taxonomic group found in natural sites (n=90), perceived areas of high species richness (HighSpcRich, n=10), plantations (n=15) arable land (n=15), and plantations (n=15). Gallers/miners represent multiple insect taxa (see appendix B for a full list). Data for insects are given per order with additional rows for the species-rich families of Carabidae and Syrphidae. The number of unique species for each habitat type and taxonomic group is given in brackets. Across sites coverage is the proportion of species likely to be found across all 130 sites, which were actually observed as estimated by extrapolation using the iNEXT package. Within sites coverage is the mean of the site specific coverage values across the 130 sites for invertebrates with abundance data. The number of new species for Denmark found during the project is also given for each taxonomic group.

The inventory was unprecedented in detail for Denmark and resulted in a total of 110 new macrofungi, 1 new lichen and 32 new invertebrate species (of which 12 were gallers and miners and 3 spiders) that had not previously been documented in Denmark (Table 2).

Turnover of plant communities among sites was adequately described by the NMDS ordination, which accounted for 81 % of the variation in plant species composition (when correlating the original distance matrix with distances in ordination space, 3-dimensional, final stress = 0.102) of which 26 %, 26 %, and 11 % could be attributed to axis 1, 2 and 3, respectively. Likewise for macrofungal communities the NMDS ordination accounted for 72 % of the variation in species composition (3-dimensional, final stress = 0.146) of which 35 %, 21 % and 14 % could be attributed to axis 1, 2, and 3, respectively. The major gradients in plant species composition of the 130 sites correlated strongly with soil fertility (NMDS axis 1 strong correlation with soil N, P and pH), successional stage (NMDS axis 2 strong correlation with light intensity and opposite correlation with litter mass and number of large trees) and soil moisture (NMDS axis 3 strong correlation with measured soil moisture), reflecting the gradients that the sites were selected to cover (Fig. 5, see correlation matrix for the rest of the environmental variables in Appendix H). Macrofungal species composition showed the same gradients, however succession and fertility swapped with succession as primary gradient (NMDS1) and fertility as secondary gradient (NMDS2). NMDS axis 3 reproduced a strong correlation with soil moisture.

**Figure 5:**
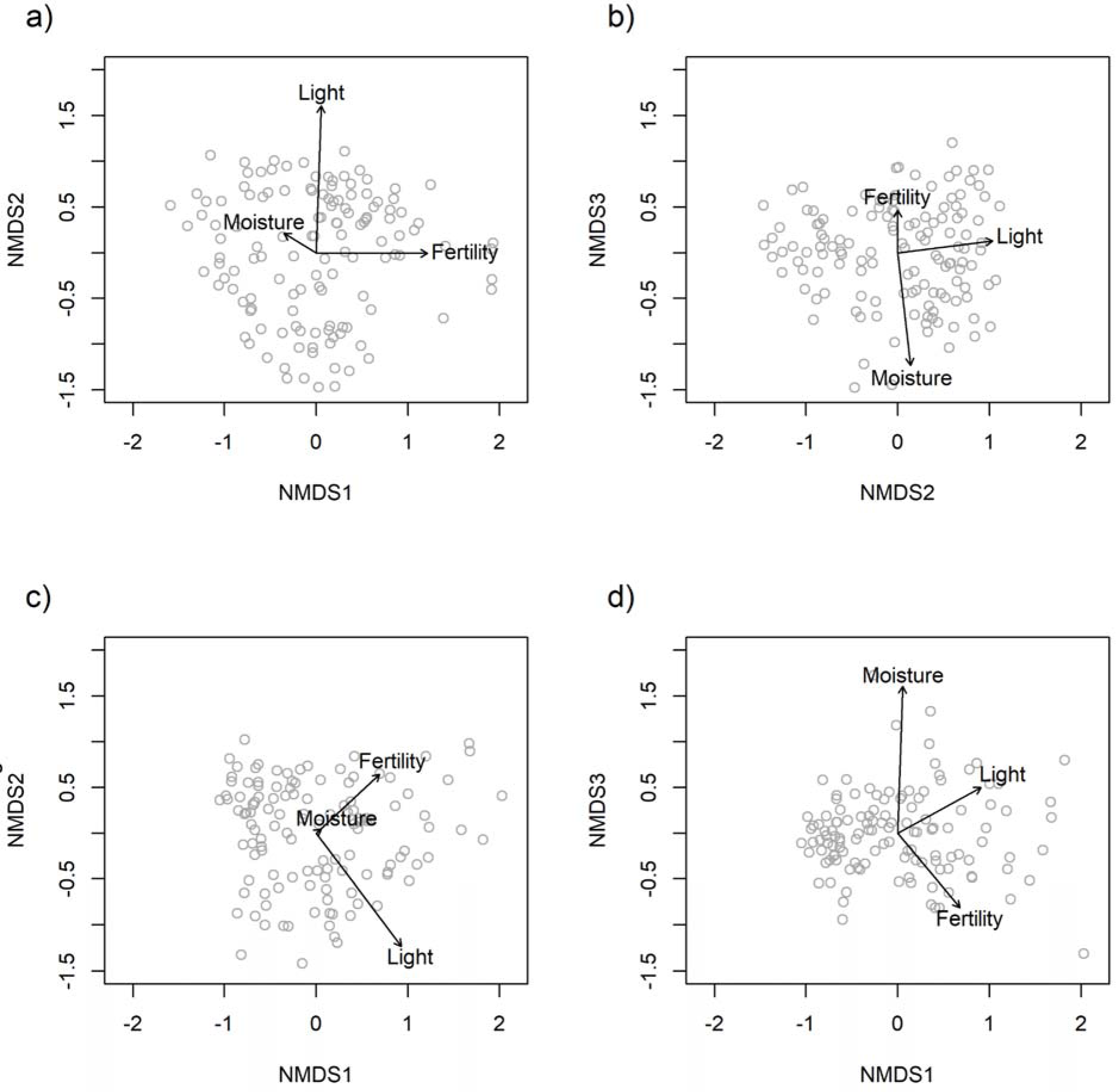
Three dimensional NMDS plots for plants with a) showing axis 2 against axis 1 and b) showing axis 3 against axis 2 and fungi with c) showing axis 2 against axis 1 and d) showing axis 3 against axis 1. The three main gradients used for selecting the 130 sites (fertility, moisture, successional stage) are overlaid as arrows (from an envfit analyses in the R package Vegan). The ordinations are based on plant species lists from the 130 sites a) & b) or macrofungi species lists from the 124 sites with more than five species c & d) and the arrows reflect soil moisture measured using a soil moisture meter, fertility measured as soil N and light measured as light intensity using HOBO loggers. The ordination plots illustrate that the community composition of vascular plants and macrofungi actually reflect the main gradients the sites were selected to cover. The scatter of dots shows the variation in abiotic conditions across the 130 sites. Correlations and p-values can be seen in Appendix H.

Spearman Rho correlations between observational plant species richness and eDNA OTU ‘richness’ as well as observational plant community composition (as represented by NMDS axes 1-3) and eDNA OTU composition were both strong and confirmative for a recovery of plant diversity by metabarcoding of soil-derived DNA (R^2^_richness_ = 0.652, R^2^_richness_ = 0.577-697, Fig. 6). Plant diversity (richness and composition) inferred from soil derived DNA thus resembled similar metrics derived from direct observation of plant communities, which has also been investigated in more detail in [40]. We found cross-correlations among species richness of different taxonomic groups to be predominantly positive or non-significant (Fig. 7). Negative correlations typically involved insect taxa like Diptera, Lepidoptera, and Orthoptera and e.g. Fungi.

**Figure 6:**
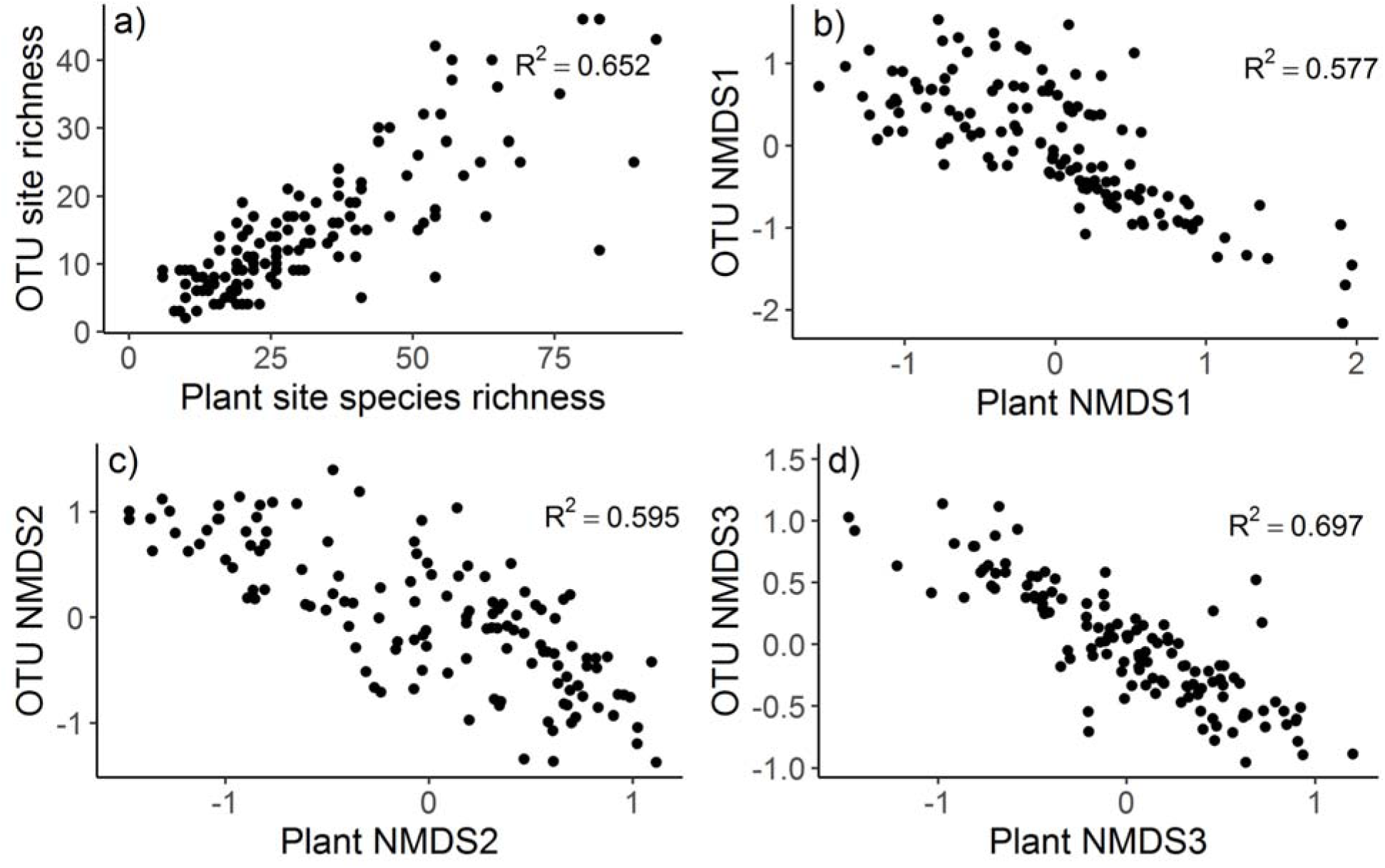
Correlation between a) observed site plant species richness and plant OTU site ‘richness’ for the 130 sites (Spearman Rho: R^2^=0.652, S = 70457, p-value < 0.001), b-d) observed site plant community composition and plant OTU community composition for the 130 sites b) NMDS axes 1 (Spearman Rho: R^2^=0.576, S = 644210, p-value < 0.001), c) NMDS axes 2 (Spearman Rho: R^2^=0.594, S = 648480, p-value < 0.001), and d) NMDS axes 3 (Spearman Rho: R^2^=0.697, S = 671850, p-value < 0.001).

**Figure 7:**
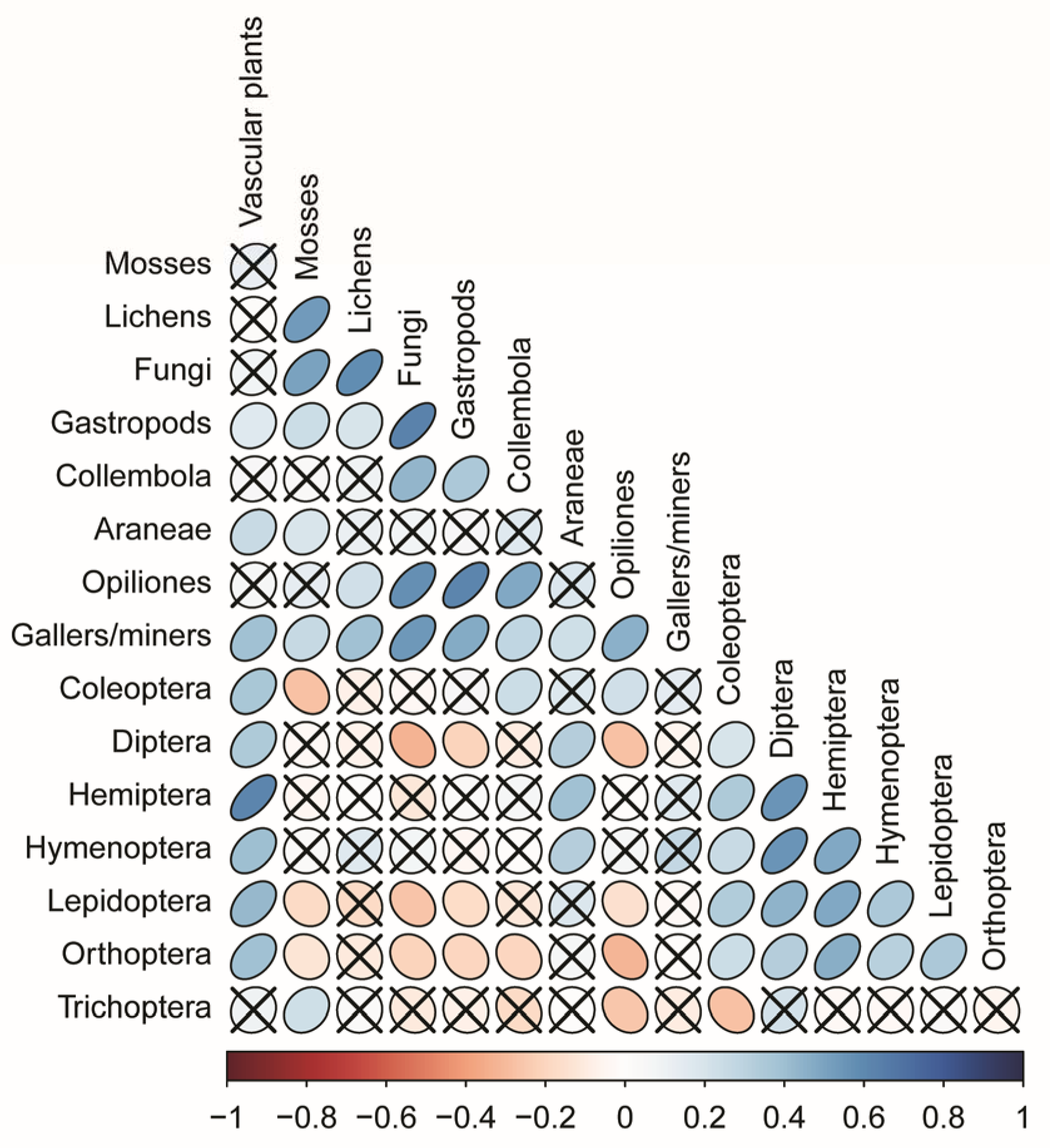
Cross correlation among the main taxonomic groups included in the study. The colour and shape of the symbol is scaled according to spearman rank correlation coefficients and non-significant (p>0.05) correlations are indicated by a cross.

## Discussion

Using ecospace as a conceptual framework [15], we developed a sampling design for mapping terrestrial biodiversity across Denmark represented by numerous, diverse taxa. Across the 130 surveyed sites, covering a tiny fraction (0.0005 %) of the total land area of Denmark, we observed approximately 5500 species, of which 143 represented new species records for the country. Our stratification procedures allowed us to cover the local and national environmental variation across Denmark using only 130 sites of 40 × 40 m each and provided a good across and within site coverage of diverse groups of invertebrates and fungi. Finally, the study demonstrates that eDNA data, once properly curated [40], can be used as an important supplement to classical biodiversity surveys.

Since, environmental filtering is an important process in community assembly [41], the most obvious design principle for a biodiversity inventory is to stratify sampling according to major abiotic and biotic environmental gradients [e.g. 42]. In strongly human-dominated landscapes, such stratification should incorporate both cultivated and non-cultivated areas and since environmental gradients are often narrower in cultivated areas, this needs to be taken into account. We found a close correspondence between the variation in average Ellenberg values at our sites and those extracted from a very large vegetation database comprising vascular plant species lists from a national monitoring program. This indicates that we managed to cover the main environmental gradients found across Denmark. Turnover of plant and macrofungi communities was significantly linked to moisture, light and fertility and allows us to generalize relationships between environment and biodiversity derived from local measurements to a large spatial extent. We note that the use of stratified random sampling implies a biased representation of rare and common environmental conditions. On the other hand, a completely random sampling would have led to limited representation of natural biotopes and their disproportionate contribution to the total biodiversity may have been missed. Our results indicate that our sampling design and site selection was successful both regarding unbiased taxonomic coverage in geographic regions and habitat types.

While the ecospace framework helped structure our sampling, it also proved challenging with respect to trade-offs between site size and homogeneity (related to ecospace position), methods to quantify biotic resources (assessing ecospace expansion) and definitions of temporal and spatial continuity. Ideally, abiotic and biotic conditions should be homogenous across a site in order to ensure that site measurements reflect the abiotic position and biotic expansion [15]. A smaller area would be more likely to be homogenous, but would be less representative. Across long environmental gradients, homogeneity and representativeness may also vary among for example, grassland, heathland, and forest. Similarly, while counting the number of different plant species is easy, accounting for the relative contribution of each species to total biomass and measuring the availability of different biotic resources such as dead wood, woody debris, litter, dung, flowers and seeds is much harder but likely to be highly important as indicated by the differences across habitat types. Finally, spatial and temporal continuity is hard to quantify due to data limitations and because past soil tillage, fertilization, or other land management or disturbance regimes have not been recorded and must be inferred indirectly. In addition, an unambiguous definition of continuity breaks is impossible given that most land use changes and derived community turnover occur gradually over time. We estimated spatial continuity using broad habitat classes at a range of scales (500 m, 1000 m, 2000 m, 5000 m) acknowledging that the dependency on spatio-temporal continuity depend on the mobility, life history and habitat specificity of different species. Our estimate of temporal continuity were also limited by the availability of aerial photographs and maps, which while not perfect, is good relative to other parts of the world. Despite these constraints, our estimates of spatial and temporal continuity varied among sites and were uncorrelated, which allowed us to statistically test for their relative roles.

We aimed at equal sampling effort per site in terms of trapping and searching time. However, this was challenged by an array of practicalities. The preferred species sampling methods varied among taxonomic groups [43, 44] and despite our application of a suite of methods, including passive sampling in pitfall traps and Malaise traps, baited traps, soil core sampling and active search, our taxonomic coverage was still incomplete (e.g. aphids, phorid flies and other species-rich groups living in the canopy are inevitably under-sampled). Our budget also forced us to be selective with the morphology-based identification of the most difficult species groups, in particular within Hymenoptera and Diptera. Among identified groups, across-site sample coverage was consistently high (>0.86) and typically close to 1, which indicates that very few unseen species remain to be recorded in each community. Invertebrate sampling and identification is extremely time consuming and relies on rare taxonomic expertise. The within site sample coverage could only be calculated for spiders and insect orders for which abundance data were available. Median values of within site sample coverage were also consistently above 0.5, which we consider adequate for cross-site comparisons. We spent more than half of the inventory budget on invertebrate sampling and identification. Invertebrates constitute by far the largest fraction of the total biota and, for many species, the adult life stage is short-lived, highly mobile, and the range of active species varies with season [45, 46]. Trapping also implies a certain risk of suboptimal placement or vandalism by visiting humans, domestic livestock or wild scavengers. The resulting number of invertebrate species per site is relatively high and revealed a considerable variation, which gives ample opportunity for comparative analyses. Although, we did not obtain full coverage of all species in every habitat category, the relative distribution of sites in arable habitats, plantations and natural sites seemed sufficient as reflected by comparable saturations of the three species accumulation curves. The high number of new species for Denmark, particularly macrofungi, can most likely be attributed to the effort, but also to the inclusion of habitat types that would otherwise have been avoided or overlooked during opportunistic field surveys [3]. Limited budgets in biodiversity studies may justify monitoring of a smaller number of taxonomic groups representing the overall biodiversity as indicated by the positive cross correlations among most taxonomic groups.

Although methods for DNA extraction, amplification, sequencing and bioinformatics processing are continuously improved and may lead to better biodiversity metrics from environmental samples, collecting representative samples from larger areas with unevenly distributed species remains a challenge. We pooled and homogenized large amounts of soil, followed by extraction of intracellular as well as extracellular DNA, from a large subsample, to maximize diversity coverage within a manageable manual workload. Biodiversity metrics based on plant DNA were correlated to the same metrics for observational plant data. This indicates that the procedure for sampling, DNA extraction and amplification can be assumed to be adequate for achieving amplicon data to quantify variation in biodiversity across wide ecological and environmental gradients for plants, but most likely also for other organisms present in the soil. These methods are promising for biodiversity studies of many organism groups that are otherwise difficult to sample and identify (e.g. nematodes, fungi, protists, and arthropods). High throughput sequencing (HTS) methods produce numerous errors [e.g. 47, 48] and it has been suggested that richness measures should be avoided altogether for HTS studies [49]. Despite the remaining challenge of relating genetic units to well-known taxonomic entities, our results along with those presented in [40] indicate that reliable metrics of α-diversity and community composition are achievable. With respect to taxonomic annotation, reference databases are far from complete and the taxonomic annotation of reference sequences are often erroneous.

Furthermore, for many groups of organisms, we have still only described and named a fraction of the actual species diversity, and the underlying genetic diversity within and between species is largely unknown for most taxa, leading to uncertainties in OTU/species delimitation and taxonomic assignment of sequence data. This also means that ecological interpretation of OTU/species assemblages assessed by eDNA is largely impossible as there is little ecological knowledge that can be linked to OTUs. Thus, for eDNA-based biodiversity assessment to further mature, molecular biologists, ecologists, and taxonomists need to work closely together to produce well-annotated reference databases. Our environmental samples for eDNA, including soil and litter samples as well as extracted DNA will be preserved for the future. This material represents a unique resource for the further development of methods within ecology and eDNA. As more efficient technologies become available in the future, it will be possible to process this material at an affordable cost and derive further insights on the relationship between traditional species occurrence, OTU data and environmental variation.

## Conclusion

We have presented a comprehensive sampling design to obtain a representative, unbiased sample of multi-taxon biodiversity stratified with respect to the major abiotic gradients. By testing and evaluating the sampling design, we conclude that it is operational and that observed biodiversity variation may be attributed to measured abiotic and biotic variables. We developed our sampling design based on the ecospace concept, and with this study, we took the first step towards general models and model inferences with transferability to terrestrial ecosystems and biotas in other parts of the world. Given the overall extent of the environmental gradients remains constant through time we believe the sampling design is also useful for monitoring biodiversity i.e. tracking changes in biodiversity through time. Meta-barcoding of environmental DNA offers a promising supplement to traditional inventories (economically and logistically), but barcode reference libraries are still far from complete. Thus, combining classical taxonomic identification with metabarcoding of environmental DNA currently appears to offer a promising approach to biodiversity research.

## Additional files

**Additional file 1: Appendix A:** Site characteristics for each of the 130 40 × 40 m sites.

**Additional file 2: Appendix B:** Sampling design for data collection.

**Additional file 3: Appendix C:** Ranges of environmental (abiotic and biotic) variables measured within the 130 sites as well as species richness of various taxonomic groups.

**Additional file 4: Appendix D:** Temporal and spatial continuity for the 130 Biowide sites.

**Additional file 5: Appendix E:** Relative richness of arthropods, bryophytes, gastropods, lichens, macrofungi and vascular plants across all 130 sites

**Additional file 6: Appendix F:** Number of species in each arthropod family for natural habitats, perceived areas of high species richness, arable land and plantations.

**Additional file 7: Appendix G:** Species accumulation curves for arable sites, plantations and natural sites

**Additional file 8: Appendix H:** Correlation matrix for NMDS axes 1, 2 and 3 and environmental variables

## Declarations

### Ethics approval and consent to participate

Not applicable

### Consent for publication

Not applicable

### Availability of data and material

The datasets used and/or analysed during the current study are available from the corresponding author on reasonable request.

### Authors’ contributions

AKB, HHB, AC, TGF, TL, AJH, MDDH and RE conceived and designed the study. AKB, HHB, LB, KF, TGF, IG, TL, GN, LS, US, and RE conducted field work. AKB, RE, LD, TTH, and TGF analyzed the data and prepared the figures. LB, KF, IG, MDDH, TL, LS, US, AAI, and HHB sorted and identified specimens. AKB, HHB, LB, ATC, KF, IG, MDDH, TTH, TGF, TL, GSN, LS, US, and RE wrote the manuscript. All authors have read and approved the final version of the manuscript.

## Acknowledgements

We thank Ako O. Mirza for plant and soil lab work, Vagn Alstrup and Roar Skovlund Poulsen for lichen surveys, volunteers that have helped in data collection, land owners, Karl-Henrik Larsson for aid in identifying critical corticioid fungi, Leif Örstadius for identifying Psathyrella collections. In regard to identification of invertebrate specimens, we would like to thank Henning Petersen for identifying Springtails (Collembola), Hjalte Kjærby for identifying Grasshoppers (Orthoptera) and Harvestmen (Opilliones), Kåre Fog for identifying snails (Gastropoda), Kåre Würtz Sørensen for identifying various Wasps (Symphyta, Spechidae, Crabronidae, and Vespidae), Lars Dyhrberg Bruun for identifying spiders (Araneae), Lars Brøndum for identifying Hoverflies (Syrphidae), Carrion Beetles (Silphidae) as well as Scarabs (Scarabidae), Lars Skipper for identifying True bugs (Heteroptera), Maja Møholt for identifying Dung beetles (Aphodius, Onthophagus and Geotrupidae) and Cantharidae, Marianne Graversen for identifying Longhorn beetles (Cerambycidae) and Ladybugs (Coccinellidae), Peter Wiberg-Larsen for identifying Caddisflies (Trichoptera), Mathias Holm for identifying True weevils and Seed weevils (Curculionidae and Apionidae), Monica Aimeé Harlund Oyre for identifying various Dipterans (Syrphida, Tachinadae, Stratiomyidae, Acroceridae, Rhagionidae, Tephritidae, Plastytomatidae, Asilidae) as well as Strepsipterans (Strepsiptera) and Book- and Barklice (Psocoptera), Morten D. D. Hansen for identifying Bees (Apoidea), Carrion beetles (Silphidae), Click beetles (Elateridae), Scarabs (Scarabaeidea) and Dung beetles (*Aphodius*, *Onthophagus* and Geotrupidae), Ole Fogh Nielsen for identifying net-winged insects (Neuroptera) and Strepsipterans (Strepsiptera), Oskar Liset Pryds Hansen and Emil Skovgaard Brandtoft for identifying Ground beetles (Carabidae), Sofie Amund Kjeldgaard and Steffen Kjeldgaard for identifying Owlet moths (Noctuidae), Mathias G. Skytte for identifying Rove beetles (Staphyllinidae), Simon Haarder for identifying galling and mining arthropods, and Ulrik Hasle Nielsen for identifying Cicadas (Cicadoidea) as well as numerous other volunteers. In regard to carrying out the eDNA lab work we would like to thank Anne Aagaard Lauridsen, Sarah Mak, Stine Raith Richter, Carlotta Pietroni and Ida Broman Nielsen. Thomas Stjernegaard Jeppesen (GBIF) is thanked for making the fungal eDNA dataset publicly available through the GBIF portal.

