## Supplemental material for "A systematic survey of regional multitaxon biodiversity: evaluating strategies and coverage"

**Appendix**

**Appendix A:** Table holding the information about class and UTM coordinates for each of the 130 sites included in the survey. The natural sites (n = 90) were selected to cover three gradients (successional stage, moisture, fertility) and the class for these were the combination of the *a priori* assumed levels of the three gradients successional stage, moisture, and fertility. The sites representing the cultivated landscape (n =30) were grouped in six classes of which three were plantations (beech, oak, and spruce) and three were arable (rotational, oldfield, ley), and finally 10 sites were assigned to the class perceived areas of high species richness (highspcrich). The table also provides the site ID, assumed successional stage (early, mid, late), assumed moisture level (dry, moist, wet), and assumed fertility level (poor, rich) for the natural sites as well as the coordinates (UTMX and UTMY).

| Site ID | | Class | | Successional stage | | Moisture | | Fertility | | UTMX | | UTMY |
| --- | --- | --- | --- | --- | --- | --- | --- | --- | --- | --- | --- | --- |
| 1 | EarlyDryPoor | | Early | | Dry | | Poor | | 581439 | | 6388139 | |
| 2 | EarlyWetRich | | Early | | Moist | | Rich | | 564449 | | 6383379 | |
| 3 | EarlyWetPoor | | Early | | Wet | | Poor | | 583805 | | 6390025 | |
| 4 | Rotational | | - | | - | | - | | 555479 | | 6375310 | |
| 5 | HighSpcRich | | - | | - | | - | | 552708 | | 6375363 | |
| 6 | Ley | | - | | - | | - | | 564600 | | 6382577 | |
| 7 | MidMoistRich | | Mid | | Moist | | Rich | | 563198 | | 6383594 | |
| 8 | MidWetPoor | | Mid | | Wet | | Poor | | 583990 | | 6389854 | |
| 9 | Spruce | | - | | - | | - | | 574170 | | 6384473 | |
| 10 | EarlyDryRich | | Early | | Dry | | Rich | | 479150 | | 6330451 | |
| 11 | EarlyMoistPoor | | Early | | Moist | | Poor | | 497046 | | 6333482 | |
| 12 | LateMoistPoor | | Late | | Moist | | Poor | | 473347 | | 6321637 | |
| 13 | LateWetPoor | | Late | | Wet | | Poor | | 495753 | | 6323436 | |
| 14 | MidDryPoor | | Mid | | Dry | | Poor | | 480429 | | 6330041 | |
| 15 | MidMoistPoor | | Mid | | Moist | | Poor | | 478318 | | 6324607 | |
| 16 | Oldfield | | - | | - | | - | | 494962 | | 6323037 | |
| 17 | Oak | | - | | - | | - | | 494653 | | 6322392 | |
| 18 | EarlyMoistRich | | Early | | Moist | | Rich | | 492405 | | 6322388 | |
| 19 | HighSpcRich | | - | | - | | - | | 552576 | | 6301441 | |
| 20 | LateDryPoor | | Late | | Dry | | Poor | | 551028 | | 6297143 | |
| 21 | LateDryRich | | Late | | Dry | | Rich | | 556187 | | 6305594 | |
| 22 | LateMoistRich | | Late | | Moist | | Rich | | 574000 | | 6311083 | |
| 23 | LateWetRich | | Late | | Wet | | Rich | | 574846 | | 6310504 | |
| 24 | MidDryRich | | Mid | | Dry | | Rich | | 548846 | | 6298056 | |
| 25 | MidWetRich | | Mid | | Wet | | Rich | | 550047 | | 6299093 | |
| 26 | Beech | | - | | - | | - | | 551954 | | 6296985 | |
| 27 | EarlyDryPoor | | Early | | Dry | | Poor | | 446101 | | 6241311 | |
| 28 | EarlyWetPoor | | Early | | Wet | | Poor | | 461377 | | 6224901 | |
| 29 | Rotational | | - | | - | | - | | 450191 | | 6239198 | |
| 30 | MidDryPoor | | Mid | | Dry | | Poor | | 469509 | | 6233884 | |
| 31 | MidMoistRich | | Mid | | Moist | | Rich | | 462049 | | 6224867 | |
| 32 | MidWetPoor | | Mid | | Wet | | Poor | | 461031 | | 6225250 | |
| 33 | MidWetRich | | Mid | | Wet | | Rich | | 448189 | | 6241792 | |
| 34 | Oldfield | | - | | - | | - | | 446448 | | 6231765 | |
| 35 | Beech | | - | | - | | - | | 468639 | | 6233492 | |
| 36 | EarlyMoistRich | | Early | | Moist | | Rich | | 448267 | | 6186362 | |
| 37 | EarlyWetRich | | Early | | Wet | | Rich | | 448741 | | 6181179 | |
| 38 | HighSpcRich | | - | | - | | - | | 452316 | | 6170040 | |
| 39 | LateMoistPoor | | Late | | Moist | | Poor | | 449133 | | 6172346 | |
| 40 | Ley | | - | | - | | - | | 450647 | | 6183553 | |
| 41 | MidDryRich | | Mid | | Dry | | Rich | | 441993 | | 6157820 | |
| 42 | MidMoistPoor | | Mid | | Moist | | Poor | | 447794 | | 6171266 | |
| 43 | Oak | | - | | - | | - | | 450717 | | 6169923 | |
| 44 | EarlyDryRich | | Early | | Dry | | Rich | | 493481 | | 6110733 | |
| 45 | EarlyMoistPoor | | Early | | Moist | | Poor | | 495874 | | 6114067 | |
| 46 | HighSpcRich | | - | | - | | - | | 497380 | | 6096051 | |
| 47 | LateDryPoor | | Late | | Dry | | Poor | | 496882 | | 6117838 | |
| 48 | LateDryRich | | Late | | Dry | | Rich | | 503771 | | 6127399 | |
| 49 | LateMoistRich | | Late | | Moist | | Rich | | 498339 | | 6096492 | |
| 50 | LateWetPoor | | Late | | Wet | | Poor | | 499012 | | 6096769 | |
| 51 | LateWetRich | | Late | | Wet | | Rich | | 504292 | | 6127348 | |
| 52 | Spruce | | - | | - | | - | | 496808 | | 6115356 | |
| 53 | EarlyDryRich | | Early | | Dry | | Rich | | 595014 | | 6228997 | |
| 54 | Rotational | | - | | - | | - | | 591617 | | 6239515 | |
| 55 | HighSpcRich | | - | | - | | - | | 614408 | | 6240840 | |
| 56 | LateDryRich | | Late | | Dry | | Rich | | 591098 | | 6239031 | |
| 57 | MidDryPoor | | Mid | | Dry | | Poor | | 594494 | | 6230173 | |
| 58 | MidDryRich | | Mid | | Dry | | Rich | | 595138 | | 6218271 | |
| 59 | MidMoistPoor | | Mid | | Moist | | Poor | | 594188 | | 6217924 | |
| 60 | MidMoistRich | | Mid | | Moist | | Rich | | 598418 | | 6231974 | |
| 61 | EarlyWetPoor | | Early | | Wet | | Poor | | 596888 | | 6232144 | |
| 62 | EarlyDryPoor | | Early | | Dry | | Poor | | 525829 | | 6215685 | |
| 63 | MidWetPoor | | Mid | | Wet | | Poor | | 522324 | | 6217641 | |
| 64 | EarlyMoistRich | | Early | | Moist | | Rich | | 543873 | | 6212230 | |
| 65 | EarlyMoistPoor | | Early | | Moist | | Poor | | 524781 | | 6215998 | |
| 66 | EarlyWetRich | | Early | | Wet | | Rich | | 545568 | | 6213415 | |
| 67 | HighSpcRich | | - | | - | | - | | 539737 | | 6214622 | |
| 68 | LateDryPoor | | Late | | Dry | | Poor | | 538527 | | 6224868 | |
| 69 | LateMoistPoor | | Late | | Moist | | Poor | | 532798 | | 6221134 | |
| 70 | LateWetPoor | | Late | | Wet | | Poor | | 530849 | | 6220234 | |
| 71 | LateMoistRich | | Late | | Moist | | Rich | | 553506 | | 6170326 | |
| 72 | LateWetRich | | Late | | Wet | | Rich | | 545923 | | 6172033 | |
| 73 | Ley | | - | | - | | - | | 543418 | | 6162941 | |
| 74 | MidWetRich | | Mid | | Wet | | Rich | | 543671 | | 6162515 | |
| 75 | Oldfield | | - | | - | | - | | 534312 | | 6169169 | |
| 76 | Beech | | - | | - | | - | | 553916 | | 6171233 | |
| 77 | Spruce | | - | | - | | - | | 554670 | | 6171805 | |
| 78 | Oak | | - | | - | | - | | 547299 | | 6171972 | |
| 79 | EarlyDryPoor | | Early | | Dry | | Poor | | 686535 | | 6212641 | |
| 80 | HighSpcRich | | - | | - | | - | | 686452 | | 6212178 | |
| 81 | LateDryPoor | | Late | | Dry | | Poor | | 704258 | | 6206120 | |
| 82 | LateMoistRich | | Late | | Moist | | Rich | | 704170 | | 6207381 | |
| 83 | LateWetPoor | | Late | | Wet | | Poor | | 706780 | | 6211031 | |
| 84 | MidWetPoor | | Mid | | Wet | | Poor | | 705403 | | 6209261 | |
| 85 | MidWetRich | | Mid | | Wet | | Rich | | 693213 | | 6212931 | |
| 86 | Spruce | | - | | - | | - | | 691428 | | 6215259 | |
| 87 | Oak | | - | | - | | - | | 702774 | | 6209432 | |
| 88 | EarlyDryRich | | Early | | Dry | | Rich | | 652603 | | 6189462 | |
| 89 | EarlyMoistPoor | | Early | | Moist | | Poor | | 643345 | | 6179060 | |
| 90 | EarlyWetPoor | | Early | | Wet | | Poor | | 642881 | | 6178095 | |
| 91 | EarlyMoistRich | | Early | | Moist | | Rich | | 638344 | | 6175468 | |
| 92 | EarlyWetRich | | Early | | Wet | | Rich | | 640495 | | 6175364 | |
| 93 | MidDryPoor | | Mid | | Dry | | Poor | | 651753 | | 6190536 | |
| 94 | MidDryRich | | Mid | | Dry | | Rich | | 618894 | | 6178182 | |
| 95 | Oldfield | | - | | - | | - | | 619112 | | 6178861 | |
| 96 | HighSpcRich | | - | | - | | - | | 675264 | | 6155726 | |
| 97 | LateDryRich | | Late | | Dry | | Rich | | 661990 | | 6140047 | |
| 98 | Beech | | - | | - | | - | | 664334 | | 6140553 | |
| 99 | LateMoistPoor | | Late | | Moist | | Poor | | 681900 | | 6161530 | |
| 100 | MidMoistPoor | | Mid | | Moist | | Poor | | 681902 | | 6160067 | |
| 101 | LateWetRich | | Late | | Wet | | Rich | | 661358 | | 6139741 | |
| 102 | Ley | | - | | - | | - | | 660682 | | 6140028 | |
| 103 | Rotational | | - | | - | | - | | 660837 | | 6132878 | |
| 104 | MidMoistRich | | Mid | | Moist | | Rich | | 664168 | | 6142266 | |
| 105 | LateDryPoor | | Late | | Dry | | Poor | | 579564 | | 6109181 | |
| 106 | EarlyDryRich | | Early | | Dry | | Rich | | 579489 | | 6109958 | |
| 107 | Oak | | - | | - | | - | | 579910 | | 6110352 | |
| 108 | EarlyWetPoor | | Early | | Wet | | Poor | | 583997 | | 6109030 | |
| 109 | MidDryPoor | | Mid | | Dry | | Poor | | 595693 | | 6107075 | |
| 110 | LateMoistPoor | | Late | | Moist | | Poor | | 587285 | | 6110023 | |
| 111 | MidMoistPoor | | Mid | | Moist | | Poor | | 587265 | | 6110673 | |
| 112 | MidWetPoor | | Mid | | Wet | | Poor | | 592504 | | 6112615 | |
| 113 | Spruce | | - | | - | | - | | 680392 | | 6073974 | |
| 114 | HighSpcRich | | - | | - | | - | | 680771 | | 6074624 | |
| 115 | Rotational | | - | | - | | - | | 660056 | | 6070320 | |
| 116 | LateWetPoor | | Late | | Wet | | Poor | | 663386 | | 6067126 | |
| 117 | LateWetRich | | Late | | Wet | | Rich | | 664394 | | 6070180 | |
| 118 | MidWetRich | | Mid | | Wet | | Rich | | 664221 | | 6069485 | |
| 119 | MidMoistRich | | Mid | | Moist | | Rich | | 670416 | | 6066245 | |
| 120 | Oldfield | | - | | - | | - | | 667308 | | 6069260 | |
| 121 | LateMoistRich | | Late | | Moist | | Rich | | 726446 | | 6098351 | |
| 122 | EarlyDryPoor | | Early | | Dry | | Poor | | 708859 | | 6105766 | |
| 123 | EarlyMoistPoor | | Early | | Moist | | Poor | | 708942 | | 6104123 | |
| 124 | EarlyMoistRich | | Early | | Moist | | Rich | | 720915 | | 6096629 | |
| 125 | EarlyWetRich | | Early | | Wet | | Rich | | 721199 | | 6095874 | |
| 126 | MidDryRich | | Mid | | Dry | | Rich | | 724661 | | 6096492 | |
| 127 | LateDryRich | | Late | | Dry | | Rich | | 726525 | | 6096762 | |
| 128 | HighSpcRich | | - | | - | | - | | 725620 | | 6098928 | |
| 129 | Ley | | - | | - | | - | | 710898 | | 6102472 | |
| 130 | Beech | | - | | - | | - | | 726483 | | 6097507 | |

**Appendix B:** **Sampling design for data collection**

**Establishing the Biowide sites**

Each Biowide site were comprised of a 40 × 40 m square, which was subdivided into four 20 × 20 m squares. Each corner of the main 40 × 40 m square was marked with at color coded flag to ensure easy identification of each sub-square during fieldwork. The colors chosen to identify the individual sub-squares were blue, red, green and yellow arranged in the indicated order clockwise around the site.

The middle of the main square was initially marked with a bamboo stick with marking tape on it but due to difficulties finding the bamboo stick within the site during fieldwork it was subsequently replaced with a white flag. The middle of each sub-square was marked with a midway-pole color coded to coincide with the color of the corresponding corner flag. During the second year of the project the flags and midway-poles in sites with grazing animals were replaced with color coded flexible plastic rods due to the animals breaking the original markings.

In order to establish the position of flags and midway-poles within each site two measuring tapes were used (one of 50 m and one of 56.6 m). Starting with the blue corner a blue flag was hammered into the ground. From this a 40 m long, straight line was measured at which point the red flag was hammered into the ground to indicate the red corner of the main square. In a 90^o^ angle from the line connecting the blue and red flag to each other a new 40 m long line was measured from the blue flag to the point where the yellow flag should be. To ensure the correct position of the yellow flag position the extended measuring tape was dragged from the red flag across the diagonal of the square to the length of approximately 56.6 m and the yellow flag was placed where the two measuring tapes intersected. At this point the center of the main square could be determined as the halfway point of the diagonal, approximately 28.3 m along the diagonal connecting the red and yellow flags. The position of the green flag was then determined by measuring a 40 m long straight line going from the yellow flag in a 90^o^ angle from the line connecting the blue and yellow flag. To ensure the correct position of the green flag the 56.6 m measuring tape was used, using the center marking as a pivot point between the red and green corner for a total length of 56.6 m. The green flag was placed where the two measuring tapes intersected. The placement of each midway-pole could then be determined as the halfway point, 14.15 m, along the line connecting each flag with the central marking of the main square.

**Sampling of plants and bryophytes**

In 2014 a list of vascular plants and epigeic bryophytes (species growing on soil) occurring within 0.5 × 0.5 m of the four plot centers was produced. A supplementary list of species within a 5 m diameter circle around the frame and not previously found in the 0.5 × 0.5 m square was produced for each of the four plots.

In 2015-2016 a supplemental survey was conducted to produce a total plant and bryophyte species list for the 130 sites. Supplemental species within the site not previously recorded within the four plots were added to the list. A list of supplemental bryophyte species was produced based on a careful examination of soil, dead wood, bark of living trees in up to 2 m height and of stones. A maximum of one hour was used on each site. Bryophyte specimens that were not possible to identify with certainty in the field were sampled and subsequently identified in the laboratory. For each bryophyte species, the substrate, e.g. phorophyte (host) species, was recorded. All records were entered in naturbasen (www. naturbasen.dk), and the nomenclature used is in accordance with this database.

**Sampling of macrofungi**

Each site was visited twice during the main fruiting season in 2014 – August-primo November – and once during the main fruiting season in 2015 - ultimo August-October either by Thomas Læssøe (TL) or TL accompanied by a volunteer – a few sites were investigated by Jacob Heilmann-Clausen. At each visit, the site was covered either clockwise or counterclockwise, and in cases with very dense herbaceous vegetation, regular inspections were carried out in kneeling position. Most woody debris was turned over to locate e.g. corticioid fungi, but there was no attempt to find hypogeous fungi, although a few were found by chance. Kneeling/crawling was used when appropriate. A visit lasted approximately 1 hour, in very open monotonous sites sometimes less, e.g. newly ploughed sites. All fruitbodies believed to belong to unique taxonomic units were sampled in one tube/site with alcohol for later sequencing (only for 2014 records). Some groups, especially corticioids and inoperculate discomycetes were mostly sampled this way and not kept for further identification. Fruitbodies of corticioid fungi were diligently collected in the 2015 campaign – excluding tomentelloid fungi - and later investigated in the lab either in living condition or as dried voucher material. Critical material was send to an expert (Karl-Henrik Larsson) for further identification. Many samples were taken back to the mobile lab for immediate microscopic investigation, and more interesting or critical material was dried as voucher material and in part deposited at the fungarium at the Natural History Museum of Denmark (herbarium C). Some critical specimens, besides the corticioids, were forwarded to external experts.

***Taxonomic coverage***

All sexual stages of Basidiomycota that form fruiting bodies were included in the sampling but the coverage of obscure jelly fungi and e.g. tomentelloid fungi cannot be considered complete. Also the sampling of Cortinarius subg. Telamonia taxa can be considered incomplete due to taxonomic difficulties. Also other members of the Cortinariaceae are not fully covered. At present a batch of Hebeloma material is being identified by Henry Beker and collaborators. Other species must be assumed not to have fruited during the visits. Rusts and smuts were not covered. Within the Ascomycota stromatic pyrenomycetes were sampled and identified to species, while other pyrenomycetous fungi were excluded except a few that readily can be identified in the field. Operculate discomycetes were sampled from wood and soil but rarely from dung. Inoperculate species with apothecia larger than 1 mm were either collected and identified or only sampled in alcohol for later sequencing. Asexual Ascomycota were largely ignored. Mycetozoa were not systematically recorded but some seemingly interesting specimens were collected and identified by an expert, they are not included in the macrofungi data set. All records were registered on www.svampeatlas.dk - a publicly available database run by the Danish Mycological Society in collaboration with the University of Copenhagen and MycoKey.

**Sampling of lichens**

For each site a total list of lichen species (lichenized fungi) was produced based on a careful examination of soil, wood, stone surfaces and bark of trees up to 2 m. A maximum of one hour was used on each site by one up to ten lichenologists including Vagn Alstrup, Roar Skovlund Poulsen and/or Ulrik Søchting at three time periods (October-November 2014, February-December 2015 and March and May 2016). Specimens that were not possible to identify with certainty in the field were sampled and subsequently identified in the laboratory. For each species the substrate, e.g. phorophyte (host) species was recorded. All records were registered in www.svampeatlas.dk, and the nomenclature used is in accordance with this database.

**Sampling of arthropods**

During 2014 two rounds of collections were performed with Malaise traps, yellow Möricke pan traps and pitfall traps, while in 2015, a single inventory with dung and carrion traps was conducted.

***Malaise traps***

To collect flying insects from the site, a Malaise trap was erected in the center of the site. The Malaise trap had the following dimensions; height at the front 190 cm, height at the back 110 cm, length 165 cm and width 115 cm. At the front, a 500 mL collection bottle was filled to two thirds with 95 % ethanol and a label containing the setup data (date, trap type, site name and UTM coordinates) was added to the bottle. In the case of grazing livestock in the site, an electric fence measuring approximately 4 × 3 m was erected around the Malaise trap to prevent the animals from knocking the trap over. After seven days, the collection bottles were collected and the captured animals and remaining ethanol were transferred to a whirl bag containing a label with the collection data (date, trap type, site name and UTM coordinates) and closed. The collection data was also noted on the outside of the bag with permanent marker. Until sorting, the animals were kept in the freezer. As the ethanol was to be used for DNA analysis it was strained from the animals before sorting using a fine mesh sieve. The ethanol was transferred to a clean whirl bag containing a data label and kept in the freezer until analysis. The sieve and other equipment was rigorously cleaned with chlorine and 95 % ethanol between each sample to prevent contamination of the samples.

***Yellow Möricke pan traps***

To collect pollinators, jumping, crawling and flying animals from the site as well as animals bouncing off the Malaise trap two yellow pan traps, measuring 42.5 cm × 31 cm × 7 cm (Length × Width × Depth) were placed underneath the Malaise trap. To inhibit the decay of collected material, the pan traps were filled to two thirds with a 5 % Rodalon solution and to reduce surface tension of the liquid, a drop of dishwashing soap was added. A label containing setup data was added to each tray. After seven days, the pan traps were collected. Collected animals were strained from the collection solution using a fine mesh fishnet after which the animals were transferred to a whirl bag containing a label with the collection data using 70 % ethanol and closed. The collection data were also noted on the outside of the bag with permanent marker. Until sorting the animals were kept in the freezer.

***Pitfall traps***

To collect the epigeic fauna from the sites, a yellow pitfall trap, 10 cm in diameter and 8 cm deep, was placed within 5 meters of each midway-pole for a total of four pitfall traps within each site. Any excavated soil was placed on a piece of plastic and after placement of the traps, excess soil was removed from the site so as to not leave any trace or disturbance within the site. If the area was grazed by livestock, two backup traps were placed within the electrically fenced area, one at each end of the Malaise trap.

The traps were filled to a quarter with a 5 % Rodalon solution to inhibit decay of the collected material and a drop of dishwashing soap was added to reduce surface tension of the liquid. A label containing the setup data was places within the trap and a piece of marking tape was placed on nearby vegetation or a bamboo stick to indicate precise location of the pitfall traps within each sub-square. After seven days, the pitfalls were collected. The collected animals were strained from the collection solution using a fine mesh fishnet after which the animals were transferred to a whirl bag containing a label with the collection data using 70 % ethanol and closed. The collection data were also noted on the outside of the bag with permanent marker. Until sorting the animals were kept in the freezer.

***Dung traps***

To collect insects associated with dung, a single collection round was performed during 2015. Fresh cow dung was collected from a herd of organic, non-treated Galloway cattle at the Mols Laboratory and frozen in 200 g packages to ensure that any dung-dwelling animals were killed.

A pitfall trap covered with chicken mesh securely fastened with tent pegs was placed within 5 meters of both the blue and green midway-pole respectively. Any excavated soil was placed on a piece of plastic and after placement of the traps excess soil was removed from the site. The traps were filled to a quarter with a 5 % Rodalon solution to inhibit decay of the collected material and a drop of dishwashing soap was added to reduce surface tension of the liquid. A label containing the setup data was placed within the trap and a piece of marking tape was placed on nearby vegetation or a bamboo stick to indicate precise location of the pitfall traps within each sub-square. On the mesh covering each trap a package of thawed out dung was placed so that any animals crawling along the ground or digging through the dung would end up in the collection solution underneath. In the case of grazing animals in the site a solid metal net basket was placed over the trap in the blue sub-square and fastened into place using tent pegs. After 4 days, the traps were collected. The collected animals were strained from the collection solution using a fine mesh sieve after which the animals were transferred to a whirl bag containing a label with the collection data using 75 % ethanol and closed. The collection data were also noted on the outside of the bag with permanent marker. Until sorting the animals were kept in the freezer.

***Carrion traps***

To collect animals associated with carrion from the site a single collection round was performed during 2015. Fresh cow heart was cut up in 2 × 2 × 2 cm pieces and frozen until use.

A pitfall trap covered with chicken mesh securely fastened with tent pegs was placed within 5 meters of both the red and yellow midway-pole respectively. Any excavated dirt was placed on a piece of plastic and after placement of the traps excess dirt was removed from the site so as to not leave any trace or disturbance within the site. The traps were filled to a quarter with a 5 % Rodalon solution to inhibit decay of the collected material and a drop of dishwashing soap was added to reduce surface tension of the liquid. A label containing the setup data was places within the trap and a piece of marking tape was placed on nearby vegetation or a bamboo stick to indicate precise location of the pitfall traps within each sub-square. A piece of thawed out cow heart was fastened securely underneath the mesh covering each trap so that any animal crawling along the ground or landing on the piece of cow heart would end up going through the mesh and end up in the collection solution underneath. In the case of grazing livestock at the site, a solid metal net basket was placed over the trap in the red sub-square and fastened into place using tent pegs. After 4 days, the traps were collected. The collected animals were strained from the collection solution using a fine mesh sieve after which the animals were transferred to a whirl bag containing a label with the collection data using 75 % ethanol and closed. The collection data were also noted on the outside of the bag with permanent marker. Until sorting the animals were kept in the freezer.

A specimen of all collected arthropod was stored in the Natural History Museum in Aarhus, Denmark.

***Sweep netting and beating***

To collect arthropods on herbs, shrubs and the lower branches of trees, a single collection round was performed during 2015 using an insect net and a beating stick. The method of sweep netting and beating was not standardized. Instead as many species as possible within a few specified taxonomic groups was collected within ~ 30 minutes. The time used on netting and beating respectively varied according to the type of vegetation. The following taxonomic groups were collected: spiders (Aranaea), true bugs (Heteroptera) and certain groups of beetles (leaf beetles (Chrysomelidae), weevils and relatives (Curculionidae etc.) and longhorn beetles (Cerambycidae)). The net used for sweeping was a cotton net (fine-meshed with a diameter of approximately 35 cm, and volume of 0.1 m^2^). The net was swiped hard against the vegetation for a high yield. Beating was conducted with a wooden stick beaten against branches of shrubs and trees as well as big and sturdy herbs, grasses etc. Simultaneously, the net was placed just under the beaten part of the plant to collect the arthropods. The collected arthropods were kept in small plastic containers in 70% ethanol until sorting.

**Sampling of galling and mining arthropods**

For each site, a list of plant-mining and gall-inducing insects and mites was produced based on scrutiny of stems, leaves, buds, flowers, fruits and underground parts of all plants at each site. Searching was carried out in a strategic way, based on the previous plant survey and on knowledge of the appearance of symptoms caused by gall and mine producing species. The field survey was done by Hans Henrik Bruun and Simon Haarder. All field work was done in the month of June (in both 2015 and 2016), chosen as to include a maximum of species irrespective of their phenology. All records were entered in Naturbasen (www.naturbasen.dk), and the nomenclature used is in accordance with this database. The data were subsequently pruned of species from groups dominated by ‘feeding’ strategies not detected by the described sampling procedure, leaving species of the following groups, Diptera: Agromyzidae and Cecidomyiidae (Subfam. Cecidomyiinae), Hymenoptera: Cynipidae, Lepidoptera: Gracillariidae and Nepticulidae, Prostigmata: Eriophyidae and Phytoptidae.

**Sampling of land gastropods**

Gastropod species were recorded by inspecting all relevant microhabitats within a site. The largest number of species was usually found by inspecting the underside of wooden substrates (branches etc.) lying on the ground; standing trunks were also inspected for resting or foraging gastropods. The underside of stones and other objects like beer cans and mussel shells lying on the ground were also inspected. Some species were mainly searched for in the litter layer, and some were found as empty shells lying on the ground, or in holes in the ground. The leaves of all relevant types of vegetation were searched, e.g. *Urtica* or *Phragmites* stands (reeds). The most important vegetation was that of sedges (*Carex spp*.) and certain grasses. Such leaves were either inspected directly, and/or they were shaken above a plastic tray, where snails shaken off were collected. In open meadow-like habitats with no wooden substrates, the shaking of grassy leaves was the main method of search. In some localities, other types of vegetation were sampled in the same way, e.g. twigs of *Vaccinium myrtillus.* In some localities, the main emphasis was on finding slugs on the fruiting bodies of fungi. In agricultural fields and hayfields, gastropods (mainly slugs) were mainly found hiding under crop remains lying on the ground. If only juveniles were found inside the site, attempts were made to find adults in the vicinity. Otherwise, juveniles were brought home and reared in captivity to adult size. In some cases, eggs were brought home and hatched to determine the species from the hatchlings. In the present investigation, it was chosen not to make any anatomical investigations, due to limited resources. Therefore, certain species could not be determined, and others only determined with some uncertainty. Further complications are due to hybridizations (in the large *Arion* species). Determinations of the two species of *Carychium* require inspection of the interior of the shell; this was carried out. A few doubtful specimens of the genus *Candidula* were sent to a Swedish expert (Ted von Proschwitz) for anatomical determination. Freshwater gastropods were included if they were found living outside of the water (*Galba truncatula* and others). Focus was on inspecting as many different microhabitats as possible, in order to maximize the number of species found. Searching was continued as long as additional species were found. The total duration of effective search varied from 30 minutes to 200 minutes. A typical search time for a moderately diverse site would be about 100 minutes. In a few sites it was evident that it would not be possible to find any gastropods, and in such cases the search time was less than 30 minutes. All sites were investigated in the October-November of 2016 avoiding periods with frost.

**Biotope data**

***Mapping of abiotic factors***
*Quadrant scale:*

Vegetation height: measured in each of the four corners of the quadrants at a 2 m distance from the quadrant center and a site mean value was calculated.

Cover: the amount of un-vegetated cover, bryophyte and lichen cover was measured in each of the four 5 m radius circles in each quadrant and a mean cover (%) was calculated for each site.

Leaf CNP: A compound sample of fresh green shoot material was collected from all plants found along a line running from the center of each quadrant to the corresponding site corners. Subsamples were analyzed for leaf carbon, nitrogen and phosphorous content. A site mean value was calculated based on the four samples.

Live woody plants: The number of live woody plants > 3 m height (< 40 cm DBH) (including climbers) were registered within the four 5 m radius circles. Each species was registered separately - but pooled for Salix, Quercus, Rosa, Malus, cherry, domestic plum, Crataegus. Quadrant counts were replaced by site counts if feasible (e.g. < 20-25 individuals at site level). For numerous taxa with no occurrences in 5 m radius circle, a number was estimated for the site.

Soil characteristics: Soil samples (0-10 cm, 5 cm diameter) were collected randomly within the four plots within each of the 130 sites and separated each sample into organic (Oa) and mineral (A/B) soil horizons. Across all sites, a total of 664 soil samples were collected. The organic horizons were separated from the mineral horizons when both were present. The depth of each soil layer was recorded using a ruler. Where appropriate, and to obtain a non-compacted sample and enough material for analysis, Oa horizons were sampled with a 23 × 17.2 cm wooden frame prior to mineral soil sampling with a core. All samples were returned to the lab and air-dried until processing. Organic soils were dried at 60°C for 48 hours, weighed, and then ground using a rotor mill. Mineral soils (2 mm) were sieved, weighed, and a subsample dried at 105°C for 48 hours to calculate gravimetric water content. Soil bulk density was measured using the oven-dry mass, area, and depth measurements collected for each core. Each mineral soil sample was ground to a fine powder using a mortar and pestle. On each sample, soil pH was measured in a slurry of 30 ml deionized water added to approximately 10g soil. The slurry was shaken vigorously for 20 seconds, allowed to settle for 30 minutes, and then measured using a Mettler Toledo Seven Compact pH meter [1]. We analyzed a subset (n = 129) of samples for total carbon and nitrogen content (LECO elemental analyzer), total phosphorus content (H_2_SO_4_-Se digestion and colorimetric analysis), and particle size (soil hydrometer method) [1]. NIR was used to analyze each sample for total carbon, nitrogen and phosphorus concentrations. Reflectance spectra was analyzed within a range of 10000-4000 cm^-1^ with a Antaris II NIR spectrophotometer (Thermo Fisher Scientific). Each spectrum with 778 data points was derived as the average of 32 scans for each if the samples. Each soil sample was mixed prior to scanning. A partial least square regression was used to test for a correlation between the NIR data and the subset reference analyses to calculate total carbon (R^2^ = 0.8619), total nitrogen (R^2^ = 0.8050), and total phosphorus (R^2^ = 0.7294) in the soil [2].
After drying soils were classified subjectively into rough soil classes by fingering moistened soil samples supplemented with visual inspection after wet sedimentation in a glass tube. We used the soil classes: Organic (more organic matter than mineral soil), Sand (not possible to form structures), Mixed sand and clay/silt/loam (loose structures may be formed, but clearly gritty texture due to sand particles), Clay/silt/loam (strong structures may be formed and soil is smooth to the touch).

*Site scale:*

Litter samples: within each of the 130 sites four litter samples were taken within a 21 cm × 21 cm frame. Litter was defined as undecomposed organic material not attached to live vegetation and included woody debris < 20 cm DBH and < 1m. The four samples from each site were pooled, taken to the laboratory, dried (60° for 48 hours) and mass (g/m^-2^) was registered.

Climate data: relative humidity (RH), light intensity, air temperature and surface temperature were measured using two HOBO loggers per site. One HOBO U23 Pro v2 Temperature/Relative Humidity Data Logger was installed 20-30 cm above ground and one HOBO Pendant^®^ Temperature/Light 8K Data Logger was installed at the ground. RH, light intensity, air temperature and surface temperature were logged once every hour during at least three weeks in the period May-August 2015 (in all 130 sites). Supplementary data for sites where loggers were out of order or vandalized by grazing animals were taken in May-August 2016 (in 37 sites). VPD (kPa) was calculated from air temperature and relative humidity using the following formula from HG Jones [3] and K Tu [4]:

$e_{s}=0.61121\times e^{17.502\times T/(240.97+T)}$ (kPa)

$e_{a}=e_{s}(\frac{RH}{100})$ (kPa)

$$VPD=e_{s}-e_{a}$$

where e_s_ is the saturated vapour pressure, e_a_ the actual vapour pressure, T the air temperature in °C, and RH the relative humidity (%).

Soil moisture: soil moisture was measured using a [FieldScout TDR 300 Soil Moisture Meter. Sixteen measurements were taken in each 40 × 40 m site](http://www.specmeters.com/soil-and-water/soil-moisture/fieldscout-tdr-meters/tdr300/) in May 2016 (spring/wet period). To cover the temporal variation in moisture the measurements were repeated in August 2016 (summer/dry period).

Boulders: we estimated the density of boulders (largest diameter > 20 cm) within each site using plotless sampling (see below). Boulders may intuitively belong to expansion as boulders are ‘habitats’ for species to live on (mosses and lichens, see below) but we categorize boulders as position because of their abiotic nature.

***Mapping of carbon sources***

*Plot scale*
Plotless sampling [BDAV3; 5] was used to estimate the density of abundant carbon sources within each of the four 5 m radius circles. Three distances were measured from each of the four quadrant centers (max distance = 5 m) to 1) closest individual, 2) nearest neighbor and 3) second nearest neighbor - or replaced by total counts if less than three items were found within the 5 m radius circle. The following carbon sources were assessed: 1) Fine woody debris (5-20 cm diameter and > 1 m, or >20 cm and < 1m), including tree stumps. 2) Flowers of insect-pollinated plants presenting flowers, irrespective of species [estimated three times (June, August 2014, April 2015]. 3) Dung of large herbivores (hare, deer, cow, horse, sheep) measured separately for each mammal species. 4) Ant hills (taller than 10 cm). 5) Small water puddles (> 0.25 m^2^, standing water, excluding short-lived (day) rain pools). Estimated density (BDAV3) for each variable was calculated at site level using the basic distance (BD) estimators for closest individual (CI), nearest neighbor (NN) and second nearest neighbor (2NN) as follows:

*BDCI = 1/(4 [∑ R_(1)i_/N]^2^)*

*BDNN = 1/(2.778 [∑ H_(1)i_/N]^2^*

*BD2N = 1/(2.778 [∑ H_(2)i_/N]^2^*

*BDAV3 = (BDCI + BDNN + BD2N)/3*

Where R_(1)i_ = the distance from the i^th^ sample point to the CI; H_(1)i_ = the distance from the i^th^ CI to its NN; H_(2)i_ = the distance from the NN at the i^th^ random point to 2NN; N = the sample size (number of random sample points used to gather distance information) [5].

*Site scale*
Large trees: The number of large live trees (> 40 DBH) within the site was registered with species name, DBH (150 cm from the largest end) and tree health index [dying = more dead than alive by volume, weakened = with wounds, hollows or active fungal attacks, but more alive than dead, healthy = only with minor closed scars, otherwise intact].

Coarse woody debris: coarse woody debris (>20 cm diameter, min length 1 m) was registered using the following parameters: lying/standing × soft/hard/fresh (decomposition class), diameter (widest, excluding stem basis) × length (only the part that was > 20 cm). Length was only measured on the part of the tree that was within the site and at least 20 cm DBH.

Uprooted areas: The number of uprooted areas with exposed unvegetated soil was registered (tree estimated to > 20 cm, DBH (150 cm from the root).

Carcasses: The number of carcasses of vertebrates with species name was registered for each site.

**eDNA data
*Soil sampling***

To get an estimate of the multicellular eukaryotic soil biodiversity, soil was collected from sites and subjected to metabarcoding through DNA extraction, PCR amplification of genetic marker regions (DNA barcoding regions) and massive parallel sequencing on the Illumina platform. The soil sampling was aiming to be representative and at the same time feasible. For each of the 130 sites several soil cores were pooled and mixed, and a subsample was used for DNA extraction.

For each site, one soil core was taken at each of the internal 81 intersections a virtual grid formed by 9 horizontal and 9 vertical lines. Sampling was carried out by pacing out this virtual grid using the corner flags and internal poles for orientation. Soil cores were taken with a thistle remover gardening tool with a curved open blade (Wolf-Garten, iWNAM 2553000) making the sampling fast and easy for most of the sampled soil types. Soil cores were scraped into a barrel (CurTec 15 litre Wide neck drum, HDPE) with a clean metal-spoon. The sampling aimed at excluding coarse leaf litter, twigs, fresh leaves and major living herbaceous plants. Each core was approximately 3-4 x 15 cm. After sampling and pooling the 81 cores of a site, the barrel was closed and transported for further processing within 24 hours. Pooled soil samples from each site weighed between 5 and 20 kg depending on soil type. The homogenizing consisted of mixing with a drilling machine (HILTI Cordless Combihammer) mounted with a mixing paddle. Mixing was carried out for at least 3 minutes and until the soil appeared homogeneous. One site with very clayey soil required addition of 2 L of water before mixing. The paddle was cleaned with a brush and in two sets of water – until no visible signs of soil was left – between samples. From each homogenized sample 5 subsamples of 5-10 spoonfuls of soil were taken out and frozen. Barrels were cleaned with tap water until no visible soil was left, and subsequently sterilized by adding, shaking 1 dl 5 % bleach solution and leaving it for at least one hour, before rinsing with clean water. This sampling design allowed for the sampling and processing of 6-9 sites per day including transportation.

The soil sampling design was inspired by a method aiming for the extraction of extracellular DNA from a large starting material [14]. We chose a similar sampling design, but decided on a more rough extraction approach involving a dedicated soil DNA extraction kit, as we wanted to target also intracellular DNA from living cells. Also, we feared that the relatively mild buffer shaking process might introduce more bias compared to a rougher physical and chemical lysis, considering the large variation of our starting material (including sand, clay, sediment-like soils, etc). The overall procedures for eDNA sampling and analyses have also been used and described in [6].

***DNA extraction***

DNA was extracted from soil using the PowerMax Soil DNA Isolation kit (MOBIO, Carlsbad, CA, USA). Four grams of soil were extracted after addition of 4 mL of 1M CaCO3 suspension, inspired by M Sagova-Mareckova, L Cermak, J Novotna, K Plhackova, J Forstova and J Kopecky [7]. DNA extract was purified using the PowerClean DNA Clean-Up Kit (MOBIO, Carlsbad, CA, USA), and DNA was normalized to 1 ng/µl. The 130 samples were extracted in smaller batches with at least one negative control for each batch.

***PCR amplification***

In this study we chose to amplify the eDNA samples with taxon specific primers for plants, fungi, earth worms, nematodes, glomeromycota and eukaryotes. For plants the Internal transcribed spacer region 2 (ITS2) was amplified with the primers S2F [8] and ITS4 [9] as well as the P6 loop of the chloroplast trnL (UAA) intron [10]. For fungi the ITS2 regions was amplified using primers gITS7 [11] and ITS4. For earth worms part of the 18S region was amplified with primers ewB and ewC from F Bienert, S De Danieli, C Miquel, E Coissac, C Poillot, JJ Brun and P Taberlet [12]. For eukaryotes, the primers 18S_allshorts from M Guardiola, MJ Uriz, P Taberlet, E Coissac, OS Wangensteen and X Turon [13] were used with a slight modification of the forward primer (TTTGTCTGGTTAATTCCG) to exclude fungi. Glomeromycota were amplified with primers NS31 [14] and AML2 [15]. Nematodes were amplified with primers used in R Sapkota and M Nicolaisen [16].

PCR amplifications contained 1X AmpliTaq Gold (Life Technologies), 0.625 μM of each primer, 0.83 mg/ml bovine serum albumin (BSA) and 1,5 μL DNA extract, 1X Gold Buffer, 2.5 mM of MgCl2, 0.08 mM each of dNTPs in 24 μL reaction volume. The number of PCR cycles was optimized with qPCR. For plant ITS2 and fungal ITS2 were used an initial denaturation step of 5 minutes at 95°C, followed by 32 cycles of denaturation of 30 seconds at 95°C, 30 seconds at 55°C, 60 seconds at 72°C, and a final elongation at 72°C for 7 minutes. For plant trnL was used an initial denaturation step of 10 minutes at 95°C, followed by 40 cycles of denaturation of 30 seconds at 95°C, 30 seconds at 55°C, 30 seconds at 72°C, and a final elongation at 72°C for 7 minutes. For earth worms was used an initial denaturation step of 10 minutes at 95°C, followed by 37 cycles of denaturation of 30 seconds at 95°C, 30 seconds at 58°C, 30 seconds at 72°C, and a final elongation at 72°C for 7 minutes. For eukaryotes was used an initial denaturation step of 5 minutes at 95°C, followed by 32 cycles of denaturation of 30 seconds at 95°C, 30 seconds at 55°C, 60 seconds at 72°C, and a final elongation at 72°C for 7 minutes. For glomeromycota was used an initial denaturation step of 10 minutes at 95°C, followed by 35 cycles of denaturation of 30 seconds at 95°C, 45 seconds at 65°C, 60 seconds at 72°C, and a final elongation at 72°C for 7 minutes. For Nematodes was used an initial PCR with untagged primers (NemF+18Sr2) using an initial denaturation step of 10 minutes at 95°C, followed by 25 cycles of denaturation of 30 seconds at 95°C, 30 seconds at 53°C, 60 seconds at 72°C, and a final elongation at 72°C for 7 minutes. A volume of 1.5 µl of diluted 1:10 initial untagged PCR product was then used as template in a nested PCR using tagged primers (NF1+18Sr2) using an initial denaturation step of 10 minutes at 95°C, followed by 20 cycles of denaturation of 30 seconds at 95°C, 30 seconds at 58°C, 60 seconds at 72°C, and a final elongation at 72°C for 7 minutes.

All primer sets were designed with 80 unique tags (MID/barcodes) of 6-8 bp at the 5’ end. No primer tag was used more than once in any sequencing library and no combination of forward and reverse primer was reused in the study. Each sample was amplified three times (if possible) using different tag combinations for approximately 400 samples per marker (130 samples + controls etc).

***Sequencing***

PCR products were pooled for a total of six pools for each marker, each pool containing half of the samples from one PCR replicate. PCR pools were cleaned with MinElute or QIAquick PCR purification kit (QIAGEN GmbH). For each marker, each of the six pools were built into separate sequencing libraries. Libraries were built using the TruSeq Dna PCR-Free Library Preparation Kit (Illumina) for plant ITS2, fungal ITS2, eukaryotes, glomeromycota and nematodes pools and NEBNext reagents (E6070) (New England BioLabs, Ipswich, MA, USA) for earth worms and plant trnL pools with some modifications to the manufacturers protocols.

Before and after library building, pools were checked on an Agilent BioAnalyzer 2100 to verify the length of the products. Adapter dimers had to be removed from some pools (plant ITS2 and fungal ITS2, eukaryote 18S, glomeromycota 18S, and nematode 18S libraries) using Agencourt AMPure XP beads. Sequencing was carried out on MiSeq (Illumina Inc., San Diego, CA, USA), at the Danish national Sequencing Centre using 150 or 250 bp PE runs, with cycle numbers adjusted according to the length of the marker: 250bp PE run for plant ITS2, fungal ITS2, nematodes, glomeromycotes and 150bp PE run for eukaryotes, plant trnL (120 cycles) and earth worms (120 cycles).

***General bioinformatics***

Sequence reads were processed using custom (unpublished) scripts based primarily on bioinformatics tools included in Cutadapt [17] DADA2 [18], VSEARCH [19] and R [20] All analyses were carried out with marker-specific optimized parameters, by first running DADA2 and then subsequently cluster OTUs at level suitable for the relevant marker. Taxonomy was assigned using custom scripts based on either VSEARCH or BLASTn to match representative sequences of each OTU against either GenBank or a in the case of dedicated reference databases – e.g UNITE [21] and Maarjam database [22]. As a final step a post-clustering OTU table curation was carried out with the LULU algorithm [23] to be able to address alfa diversity of the samples more accurately.

***Bioinformatic processing of eDNA plant data for comparison with observational data***

Reads were demultiplexed using custom scripts, and reads from PCR replicates of the same samples were merged. Reads were processed with the DADA2 r-package [18] as described here (http://benjjneb.github.io/dada2/tutorial.html) with minor adjustments to accommodate our tagging scheme. Acknowledging that the selected marker for plants is known to contain intraspecific variation, the result from the DADA2 analyses were then clustered at 97% level using VSEARCH [19]. OTUs were taxonomically annotated using custom scripts for mapping against GenBank. Only OTUs assigned to vascular plant lineages with 80% or higher sequence identity were retained for further analyses. Lastly, OTUs were filtered using a post-clustering curation method [23] to further reduce the number of erroneous OTUs.

The resulting OTU tables were then subjected to a simple richness measure (OTU count) and a community composition analysis with NMDS, using the metaMDS method from Vegan as implemented in Phyloseq [24]. NMDS analyses were carried out using Bray-Curtis dissimilarity after reducing to presence/absence data, and also on abundance (read count) data after applying a Hellinger transformation. The resulting measures were then correlated with the same measures on the observational plant data using Spearman Rho correlation.

Raw data and detailed documentation of the analyses can be obtained from Tobias Guldberg Frøslev.

**Landscape data
*Temporal continuity***

Temporal continuity refers to continuity of position and expansion. Break in continuity within a site may affect populations (we are not interested in the effect on the individuals). Temporal continuity is estimated by time since major land use change within the 40 × 40 m site. Break in continuity can be defined as a crucial change in disturbance regime for the given area (i.e. a significant change in natural or anthropogenic disturbance regime). This way, continuity of a field is broken if disturbance events are not recurrent while continuity of a forest is broken if the forest is cut down. Break in continuity should not be confused with succession (more gradual change) – although the two concepts may overlap.

As different taxonomic groups may respond differently to continuity of position (abiotic conditions: light, nutrients, hydrology) and continuity of expansion (carbon sources), ideally, the two can be distinguished. Knowledge of five variables sufficiently reflects temporal continuity: ‘soil continuity’ (e.g. plowing), dead wood (cutting and storm damage), light conditions (overgrowing, storm damage, cutting and planting), nutrient status (e.g. use of fertilizer since 1950s), water/hydrology (drainage, damming, restoration).

For each site, a temporal sequence of aerial photos was inspected starting with the most recent photos. Temporal continuity is the time since the most recent apparent major land use change. Aerial photos were available for 2014, 2012, 2010, 2008, 2006, 2004, 2002, 1999, 1995, 1954, 1945 and historical maps illustrate the period 1842-1945. In addition, Flyfotoarkivet (https://www.flyfotoarkivet.dk/) holds aerial photos for most of Jutland (not Sønderjylland) in the periode 1954-1993. Royal Air Force photos (1968) were also used although photos were not available for all sites. Temporal continuity was estimated by visual inspection of changes in land use. The aerial photos were inspected sequentially from most recent to oldest photos/maps. The year in which the change was identified was noted as ‘break in continuity’.

Special cases

*Succession*

If 2014 photo shows > 20 % open/non-vegetated cover: Estimate continuity for open landscape and continuity for shrub (continuity shrub = the number of years in which shrub cover > 20 %). If temporal continuity differs between open and shrub, the highest value is used.

If 2014 photo shows < 20 % open/non-vegetated cover: Estimate continuity for shrub (continuity shrub = the number of years in which shrub cover > 20 %).

If 2014 photo is shrubby but has >80 % tree canopy cover it is defined as forest.

*Forest*

If forest map from The Danish Nature Agency exists:

- if the species is a commonly cultivated species: the year in which the trees were planted (from map) is used. Continuity for forest is then defined to begin 5 years after time of planting.
- if the species is not used in forestry (e.g. Tilia), OR the year of origin is before 1810 (start of plantation forestry), OR if the species actually occurring at the site as overstorey are indigenous tree species AND different from the species indicated on the map: then forest maps are overruled (e.g. Lindestykket in Draved forest); instead ordnance maps (målebordsblade) and maps of the Royal Danish Academy of Sciences and Letters of 1768-1805 (Videnskabernes Selskabs Kort, http://hiskis2.dk/) are used to estimate the age (no waiting period of 5 years is added).

If forest map from The Danish Nature Agency does not exist: aerial photos are used to estimate year of planting and continuity for forest is then defined to begin 5 years after time of planting.

*Open habitats*

Aerial photos and Earliest Cadastral maps from the time of land reforms (mainly 1780-1810) are used to estimate continuity.

*Intensive farmland (annual rotation and sown, fertilized leys)*

Standard value (continuity = 64 years – since 1950).

*Continuity classes:*

After having estimated the temporal continuity for all sites we reclassified the continuity into a 4-level variable: 1: <15 years, 2: 15-45 years, 3: 45-135 years, 4: >135 years.

**Spatial continuity**

Spatial continuity refers to continuity of position and expansion. Break in continuity within a site may affect populations (we are not interested in the effect on the individuals). Spatial continuity was estimated by different levels of continuity of natural areas within a set of buffer distances from the site – to account for the fact that landscape connectivity or structural connectivity/continuity will be different for different taxonomic groups. Four buffers were constructed for each site: 500 m buffer, 1000 m buffer, 2000 m buffer and 5000 m buffer. Within the four buffers we subjectively estimated the spatial continuity of the habitat type of the site by visual interpretation of aerial photographs and additional information from land mapping of woodlands, fields, grassland, heathland, meadows, salt marshes and mires. Visual interpretation was performed by three researchers that were familiar with the sites. We calibrated estimates by identifying continuity estimates showing large deviation from the overall mean (deviation = (estimated value – overall mean)/overall mean). To down weight deviations according to the overall mean we calculated a weighted deviation (weighted deviation = deviation * overall mean/100). All estimates with weighted deviation >10 % were revisited and calibrated. The final spatial continuity measure within each buffer was the overall mean of the calibrated values.

**Appendix C:** Ranges of environmental (abiotic and biotic) variables measured within the 130 sites as well as species richness of various taxonomic groups.

| Measured variable | Min value | Max value |
| --- | --- | --- |
| Litter mass from four litter samples from each site (21x21cm frame) (g/m^2^) | 0.57 | 2738.95 |
| Soil pH in 0-10 cm soil sample. Weighted for the proportion of organic matter to mineral soil | 3.94 | 8.13 |
| Soil C 0-10 cm (g/m^2^ ) | 6.26 | 21270.62 |
| Soil N 0-10 cm (g/m^2^ ) | 5.67 | 703.82 |
| Soil P 0-10 cm (g/m^2^) | 0.15 | 115.22 |
| Soil class (four classes: sand, sand-clay, organic, clay) | - | - |
| % leaf N - mean of four quadrants | 1.29 | 4.95 |
| % leaf C - mean of four quadrants | 40.55 | 47.98 |
| % leaf P - mean of four quadrants | 0.09 | 0.84 |
| Maximum surface temperature (°C) calculated for all day | 14.33 | 61.74 |
| Minimum nighttime air temperature (°C) | -3.30 | 8.42 |
| Median light intensity calculated for all day (lux) | 16.00 | 1344.00 |
| Mean daytime vapour pressure deficit (kPa) | 0.06 | 0.86 |
| Mean site Ellenberg L | 3.63 | 7.75 |
| Mean site Ellenberg F | 3.70 | 8.73 |
| Mean site Ellenberg R | 2.31 | 6.80 |
| Mean site Ellenberg N | 1.51 | 6.81 |
| Mean site Ellenberg S | 0.00 | 0.66 |
| Mean site Ellenberg T | 3.92 | 6.00 |
| Trimmed mean soil moisture (%Volumetric Water Content (VWC)) value from 16 soil moisture meter values pr. site | 1.28 | 77.72 |
| Species number of plants in site | 11.00 | 134.00 |
| Species number of bryophytes in site | 0.00 | 50.00 |
| Species number of macrofungi in site | 0.00 | 180.00 |
| Species number of lichen in site | 0.00 | 33.00 |
| Species number of carabid beetles in site | 0.00 | 21.00 |
| Species number of hoverflies in site | 0.00 | 22.00 |
| Species number of spiders in site | 7.00 | 53.00 |
| Species number of gallers and miners in site | 0.00 | 17.00 |
| Species number of gastropods in site | 0.00 | 33.00 |
| Mean herb layer vegetation height (cm, four measurements within each of the four plots) | 0.00 | 146.69 |
| Mean % bare soil (mean of % cover in each of the four 5 m-circle plots) | 0.00 | 97.00 |
| Mean % bryophyte cover (mean of % cover in each of the four 5 m-circle plots) | 0.00 | 95.00 |
| Mean % lichen cover (mean of % cover in each of the four 5 m-circle plots) | 0.00 | 54.25 |
| Volume of dead wood (m^3^/ha) | 0.00 | 8.14 |
| Estimated density of trees < 40DBH – deciduous (number/m^2^) | 0.00 | 0.49 |
| Estimated density of trees < 40DBH - coniferous (number/m^2^) | 0.00 | 0.08 |
| Number of trees > 40DBH | 0.00 | 24.00 |
| Number of uprooted trees | 0.00 | 13.00 |
| Number of carcasses | 0.00 | 0.00 |
| Basic distance abundance estimate of deer dung (number/m^2^). BADV3 from White et al. (2008) | 0.00 | 0.98 |
| Basic distance abundance estimate of hare dung (number/m^2^). BADV3 from White et al. (2008) | 0.00 | 7.07 |
| Basic distance abundance estimate of cow dung (number/m^2^). BADV3 from White et al. (2008) | 0.00 | 0.24 |
| Basic distance abundance estimate of sheep dung (number/m^2^). BADV3 from White et al. (2008) | 0.00 | 3.89 |
| Basic distance abundance estimate of horse dung (number/m^2^). BADV3 from White et al. (2008) | 0.00 | 0.27 |
| Summed dung density (number/m^2^). BADV3 from White et al. (2008) | 0.00 | 7.07 |
| Basic distance abundance estimate of water puddles > 0.25m^2^ (number/m^2^). BADV3 from White et al. (2008) | 0.00 | 0.19 |
| Basic distance abundance estimate of dead wood (diameter 5-20 cm and length >1m or diameter >20cm and length <1m) (number/m^2^). BADV3 from White et al. (2008) | 0.00 | 3.02 |
| Basic distance abundance estimate of ant hills (height > 10cm) (number/m^2^). BADV3 from White et al. (2008) | 0.00 | 0.08 |
| Basic distance abundance estimate of boulders (diameter > 20cm) (number/m^2^). BADV3 from White et al. (2008) | 0.00 | 0.16 |
| Estimated mean flower abundance within site (number/m^2^). BADV3 from White et al. (2008) | 0.00 | 1223 |
| Temporal continuity class (4 levels: 1: <15 years. 2: 15-45 years. 3: 45-135 years. 4: >135 years) | 1.00 | 4.00 |
| Spatial continuity of focal habitat type within 500 m buffer (%) | 2.00 | 93.33 |
| Spatial continuity of focal habitat type within 1000 m buffer (%) | 1.00 | 78.33 |
| Spatial continuity of focal habitat type within 2000 m buffer (%) | 0.03 | 73.33 |
| Spatial continuity of focal habitat type within 5000 m buffer (%) | 0.03 | 71.67 |

**Appendix D:** Temporal and spatial continuity for the 130 Biowide sites. Temporal continuity is represented on a 4-level ordinal scale: 1: < 15 years of continuity, 2: 15-44 years of continuity, 3: 45-135 years of continuity and 4: >135 years of continuity (or continous on the oldest available map). Spatial continuity represents the amount of similar habitat (%) as the focal site habitat within four different buffer sizes (500m, 1000m, 2000m, 5000m).

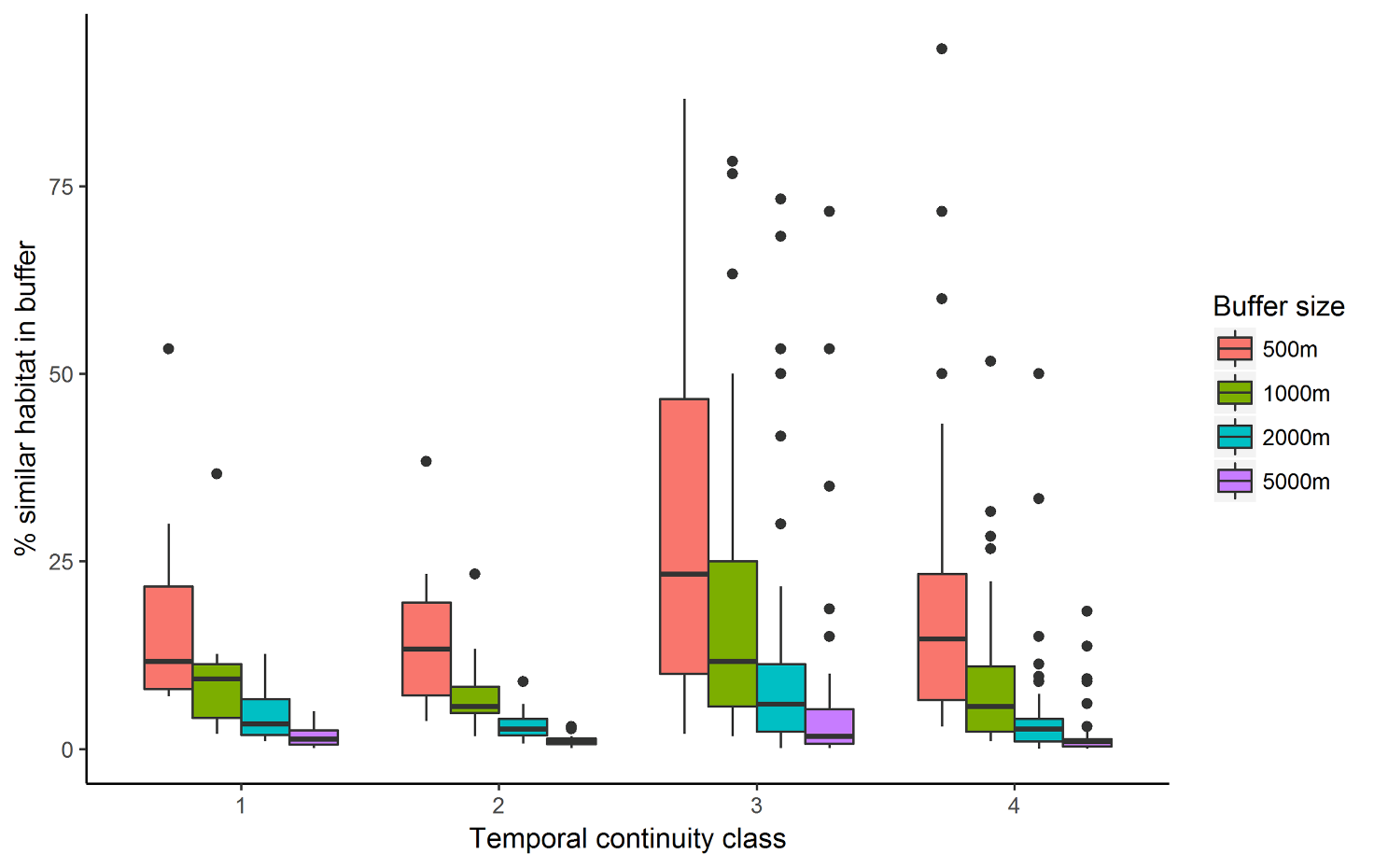

**Appendix E:** Relative richness of arthropods, bryophytes, gastropods, lichens, macrofungi and vascular plants across the five geographical regions: Eastern Jutland (Ejut), Funen, Lolland, Møn (FLM), Northern Jutland (Njut), Western Jutland (Wjut) and Zealand (Zeal). Nestedness and turnover for each region as well as average and standard deviations are given.

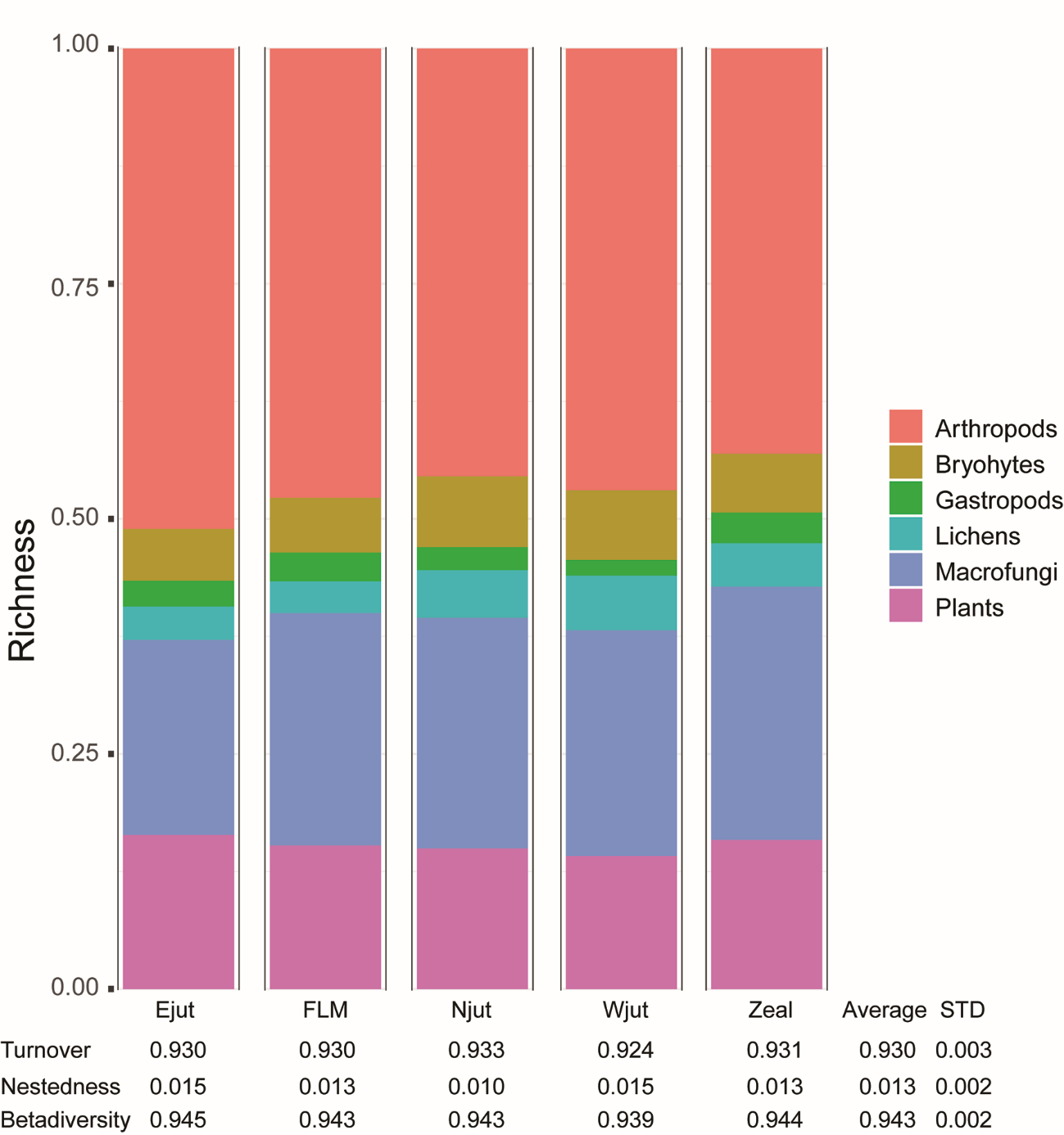

**Appendix F:** Number of species per arthropod family found in natural sites (n=90), perceived areas of high species richness (HighSpcRich, n=10), arable land (n=15), and plantations (n=15). The number of unique species for each habitat type and family is given in brackets.

| Order | Family | Total | Natural | HighSpcRich | Arable | Plantation |
| --- | --- | --- | --- | --- | --- | --- |
| Araneae | Agelenidae | 2 | 2 (0) | 1 (0) | 1 (0) | 1 (0) |
| Araneae | Amaurobiidae | 2 | 2 (0) |  |  | 2 (0) |
| Araneae | Anyphaenidae | 1 | 1 (0) | 1 (0) |  | 1 (0) |
| Araneae | Araneidae | 23 | 21 (5) | 13 (0) | 10 (0) | 8 (2) |
| Araneae | Atypidae | 1 | 1 (0) | 1 (0) |  |  |
| Araneae | Clubionidae | 13 | 13 (3) | 8 (0) | 7 (0) | 4 (0) |
| Araneae | Corinnidae | 2 | 1 (0) | 2 (1) |  |  |
| Araneae | Cybaeidae | 1 | 1 (1) |  |  |  |
| Araneae | Dictynidae | 4 | 4 (1) | 2 (0) | 2 (0) | 1 (0) |
| Araneae | Gphosidae | 17 | 17 (6) | 10 (0) | 4 (0) | 1 (0) |
| Araneae | Hahniidae | 4 | 4 (1) | 3 (0) |  | 2 (0) |
| Araneae | Linyphiidae | 144 | 132 (44) | 50 (1) | 54 (2) | 72 (8) |
| Araneae | Liocranidae | 4 | 4 (3) | 1 (0) |  | 1 (0) |
| Araneae | Lycosidae | 25 | 25 (4) | 17 (1) | 14 (0) | 3 (0) |
| Araneae | Mimetidae | 2 | 2 (1) |  | 1 (0) | 1 (0) |
| Araneae | Miturgidae | 2 | 2 (0) | 2 (0) | 2 (0) |  |
| Araneae | Oxyopidae | 1 |  | 1 (1) |  |  |
| Araneae | Philodromidae | 9 | 8 (2) | 5 (0) | 4 (0) | 4 (0) |
| Araneae | Pisauridae | 2 | 3 (1) | 1 (0) | 1 (0) |  |
| Araneae | Salticidae | 18 | 16 (11) | 3 (0) | 4 (0) | 2 (1) |
| Araneae | Segestriidae | 1 | 1 (0) |  |  | 1 (0) |
| Araneae | Sparassidae | 1 | 1 (1) |  |  |  |
| Araneae | Tetragthidae | 11 | 11 (1) | 9 (0) | 3 (0) | 8 (0) |
| Araneae | Theridiidae | 30 | 27 (11) | 12 (0) | 15 (2) | 10 (0) |
| Araneae | Theridiosomatidae | 1 | 1 (1) |  |  |  |
| Araneae | Thomisidae | 12 | 11 (5) | 5 (1) | 3 (0) | 3 (1) |
| Araneae | Uloboridae | 1 | 1 (0) |  |  | 1 (0) |
| Araneae | Zoridae | 1 | 1 (0) |  | 1 (0) | 1 (0) |
| Coleoptera | Anthribidae | 2 | 2 (1) | 1 (0) |  | 1 (0) |
| Coleoptera | Attelabidae | 7 | 5 (5) |  |  | 2 (2) |
| Coleoptera | Brentidae | 34 | 30 (13) | 8 (1) | 19 (3) | 1 (0) |
| Coleoptera | Buprestidae | 1 | 1 (1) |  |  |  |
| Coleoptera | Cantharidae | 34 | 32 (11) | 10 (0) | 10 (0) | 11 (2) |
| Coleoptera | Carabidae | 123 | 107 (43) | 34 (3) | 51 (15) | 35 (1) |
| Coleoptera | Cerambycidae | 12 | 12 (6) | 2 (0) | 1 (0) | 5 (1) |
| Coleoptera | Cleridae | 2 | 2 (2) |  |  |  |
| Coleoptera | Coccinellidae | 20 | 20 (9) | 5 (0) | 8 (0) | 5 (0) |
| Coleoptera | Curculionidae | 132 | 104 (55) | 35 (9) | 43 (13) | 32 (6) |
| Coleoptera | Dascillidae | 1 |  | 1 (1) |  |  |
| Coleoptera | Dasytidae | 2 | 2 (2) |  |  |  |
| Coleoptera | Drilidae | 1 | 1 (1) |  |  |  |
| Coleoptera | Elateridae | 22 | 21 (6) | 9 (0) | 7 (0) | 6 (1) |
| Coleoptera | Endomychidae | 1 |  |  | 1 (1) |  |
| Coleoptera | Eucnemidae | 2 | 2 (1) | 1 (0) |  | 1 (0) |
| Coleoptera | Geotrupidae | 3 | 3 (1) | 2 (0) | 2 (0) | 2 (0) |
| Coleoptera | Histeridae | 5 | 4 (2) | 2 (1) | 2 (0) |  |
| Coleoptera | Hydrophilidae | 11 | 10 (3) | 6 (1) | 5 (0) |  |
| Coleoptera | Kateretidae | 1 | 1 (1) |  |  |  |
| Coleoptera | Lampyridae | 1 | 1 (1) |  |  |  |
| Coleoptera | Latridiidae | 2 | 2 (1) |  |  | 1 (0) |
| Coleoptera | Lucanidae | 1 | 1 (1) |  |  |  |
| Coleoptera | Lycidae | 2 | 2 (0) |  |  | 2 (0) |
| Coleoptera | Malachiidae | 2 | 2 (1) |  | 1 (0) |  |
| Coleoptera | Melandryidae | 1 | 1 (1) |  |  |  |
| Coleoptera | Monotomidae | 1 | 1 (1) |  |  |  |
| Coleoptera | Nitidulidae | 2 | 2 (1) |  | 1 (0) |  |
| Coleoptera | Oedemeridae | 3 | 3 (1) | 1 (0) |  | 1 (0) |
| Coleoptera | Ptinidae | 2 | 2 (2) |  |  |  |
| Coleoptera | Pyrochroidae | 1 | 1 (1) |  |  |  |
| Coleoptera | Salpingidae | 2 | 2 (1) |  |  | 1 (0) |
| Coleoptera | Scarabaeidae | 28 | 25 (8) | 13 (1) | 17 (2) | 6 (0) |
| Coleoptera | Silphidae | 10 | 10 (0) | 7 (0) | 8 (0) | 6 (0) |
| Coleoptera | Staphylinidae | 76 | 54 (29) | 15 (6) | 25 (15) | 16 (4) |
| Coleoptera | Tenebrionidae | 3 | 3 (2) | 1 (0) | 1 (0) |  |
| Coleoptera | Throscidae | 1 | 2 (1) |  |  |  |
| Diptera | Acroceridae | 1 | 1 (0) |  | 1 (0) |  |
| Diptera | Asilidae | 8 | 7 (2) | 4 (0) | 2 (1) | 2 (0) |
| Diptera | Bombyliidae | 2 | 1 (1) | 1 (1) |  |  |
| Diptera | Cecidomyiidae | 1 | 1 (1) |  |  |  |
| Diptera | Chloropidae | 2 | 2 (2) |  |  |  |
| Diptera | Hybotidae | 1 | 1 (1) |  |  |  |
| Diptera | Micropezidae | 1 |  |  | 1 (1) |  |
| Diptera | Oestridae | 1 | 1 (1) |  |  |  |
| Diptera | Pediciidae | 1 | 1 (1) |  |  |  |
| Diptera | Platystomatidae | 2 | 1 (0) | 2 (1) | 1 (0) |  |
| Diptera | Ptychopteridae | 1 | 1 (1) |  |  |  |
| Diptera | Rhagionidae | 5 | 5 (0) | 3 (0) | 3 (0) | 5 (0) |
| Diptera | Rhinophoridae | 4 | 4 (3) | 1 (0) |  |  |
| Diptera | Scathophagidae | 1 | 1 (1) |  |  |  |
| Diptera | Sciomyzidae | 2 | 2 (2) |  |  |  |
| Diptera | Stratiomyidae | 14 | 14 (8) | 4 (1) | 6 (0) |  |
| Diptera | Syrphidae | 98 | 93 (41) | 36 (2) | 45 (6) | 20 (2) |
| Diptera | Tachinidae | 28 | 26 (14) | 7 (2) | 8 (1) | 8 (2) |
| Diptera | Tephritidae | 20 | 15 (7) | 5 (2) | 9 (3) |  |
| Diptera | Ulidiidae | 2 | 2 (2) |  |  |  |
| Diptera | Xylomyidae | 1 | 1 (1) |  |  |  |
| Hemiptera | Acanthosomatidae | 5 | 6 (3) | 1 (1) | 1 (0) | 2 (0) |
| Hemiptera | Alydidae | 1 | 1 (1) |  |  |  |
| Hemiptera | Anthocoridae | 10 | 11 (3) | 7 (1) | 5 (0) | 3 (0) |
| Hemiptera | Aphrophoridae | 5 | 5 (1) | 4 (0) | 2 (0) | 1 (0) |
| Hemiptera | Berytidae | 3 | 3 (0) | 2 (0) | 1 (0) | 1 (0) |
| Hemiptera | Caliscelidae | 1 | 1 (1) |  |  |  |
| Hemiptera | Ceratocombidae | 1 | 1 (0) | 1 (0) |  | 1 (0) |
| Hemiptera | Cercopidae | 1 | 1 (1) |  |  |  |
| Hemiptera | Cicadellidae | 158 | 169 (52) | 80 (3) | 85 (6) | 44 (2) |
| Hemiptera | Cimicidae | 1 | 1 (1) |  |  |  |
| Hemiptera | Cixiidae | 4 | 5 (2) |  | 2 (0) | 3 (0) |
| Hemiptera | Coreidae | 1 | 1 (1) |  |  |  |
| Hemiptera | Cydnidae | 3 | 3 (3) |  |  |  |
| Hemiptera | Delphacidae | 32 | 38 (20) | 11 (1) | 8 (1) | 4 (0) |
| Hemiptera | Gerridae | 1 | 1 (1) |  |  |  |
| Hemiptera | Lygaeidae | 33 | 35 (17) | 13 (2) | 5 (0) | 6 (1) |
| Hemiptera | Microphysidae | 4 | 5 (1) | 4 (0) | 2 (0) | 3 (0) |
| Hemiptera | Miridae | 113 | 112 (47) | 40 (4) | 34 (1) | 31 (4) |
| Hemiptera | bidae | 10 | 10 (1) | 9 (0) | 6 (1) | 5 (0) |
| Hemiptera | Nepidae | 1 | 1 (1) |  |  |  |
| Hemiptera | Pemphigidae | 1 | 1 (1) |  |  |  |
| Hemiptera | Pentatomidae | 15 | 16 (8) | 6 (0) | 4 (0) | 1 (0) |
| Hemiptera | Piesmatidae | 1 | 1 (0) |  | 1 (0) |  |
| Hemiptera | Psyllidae | 5 | 4 (2) | 2 (1) | 1 (0) |  |
| Hemiptera | Reduviidae | 4 | 3 (3) |  |  | 1 (1) |
| Hemiptera | Rhopalidae | 6 | 7 (2) | 3 (0) | 3 (0) |  |
| Hemiptera | Saldidae | 10 | 11 (9) |  | 2 (0) | 1 (0) |
| Hemiptera | Scutelleridae | 2 | 3 (2) |  |  |  |
| Hemiptera | Tingidae | 10 | 10 (2) | 6 (0) | 5 (0) |  |
| Hemiptera | Triozidae | 1 | 1 (0) | 1 (0) | 1 (0) |  |
| Hemiptera | Veliidae | 2 | 2 (2) |  |  |  |
| Hymenoptera | Ampulicidae | 1 |  | 1 (1) |  |  |
| Hymenoptera | Andrenidae | 5 | 2 (0) | 2 (1) | 2 (0) | 3 (1) |
| Hymenoptera | Aphelinidae | 1 | 1 (1) |  |  |  |
| Hymenoptera | Apidae | 14 | 13 (6) | 7 (1) | 5 (0) | 6 (0) |
| Hymenoptera | Argidae | 2 | 3 (2) |  |  |  |
| Hymenoptera | Bethylidae | 1 |  |  | 1 (1) |  |
| Hymenoptera | Chalcididae | 2 | 1 (1) | 1 (1) |  |  |
| Hymenoptera | Chrysididae | 2 | 2 (2) |  |  |  |
| Hymenoptera | Colletidae | 4 | 4 (1) | 2 (0) | 2 (0) |  |
| Hymenoptera | Crabronidae | 29 | 31 (18) | 11 (1) | 5 (1) | 3 (0) |
| Hymenoptera | Cynipidae | 3 | 3 (3) |  |  |  |
| Hymenoptera | Diapriidae | 1 |  |  |  | 1 (1) |
| Hymenoptera | Dryinidae | 2 | 2 (0) |  | 1 (0) | 2 (0) |
| Hymenoptera | Encyrtidae | 1 | 1 (1) |  |  |  |
| Hymenoptera | Evaniidae | 1 | 1 (0) | 1 (0) |  |  |
| Hymenoptera | Formicidae | 1 | 2 (1) |  |  |  |
| Hymenoptera | Halictidae | 12 | 11 (7) | 4 (2) | 2 (0) |  |
| Hymenoptera | Ichneumonidae | 3 | 3 (2) | 1 (0) |  |  |
| Hymenoptera | Megachilidae | 9 | 6 (5) | 3 (2) | 1 (1) |  |
| Hymenoptera | Megaspilidae | 1 | 2 (1) |  |  |  |
| Hymenoptera | Melittidae | 2 | 1 (1) |  | 1 (1) |  |
| Hymenoptera | Mutillidae | 1 | 1 (1) |  |  |  |
| Hymenoptera | Mymaridae | 1 | 1 (1) |  |  |  |
| Hymenoptera | Pamphiliidae | 1 |  |  |  | 1 (1) |
| Hymenoptera | Pompilidae | 11 | 12 (8) | 2 (1) | 2 (1) |  |
| Hymenoptera | Pteromalidae | 1 | 1 (1) |  |  |  |
| Hymenoptera | Sphecidae | 4 | 4 (3) | 1 (0) |  |  |
| Hymenoptera | Tenthredinidae | 56 | 54 (31) | 11 (4) | 14 (1) | 8 (2) |
| Hymenoptera | Tiphiidae | 1 | 1 (1) |  |  |  |
| Hymenoptera | Vespidae | 13 | 14 (6) | 5 (0) | 3 (0) | 4 (0) |
| Lepidoptera | Drepanidae | 2 | 2 (1) | 1 (0) |  |  |
| Lepidoptera | Erebidae | 4 | 4 (2) | 1 (0) |  | 1 (0) |
| Lepidoptera | Geometridae | 16 | 12 (6) | 5 (2) |  | 5 (2) |
| Lepidoptera | Hepialidae | 3 | 3 (2) | 1 (0) | 1 (0) |  |
| Lepidoptera | Hesperiidae | 3 | 3 (3) |  |  |  |
| Lepidoptera | Lasiocampidae | 1 | 1 (1) |  |  |  |
| Lepidoptera | Limacodidae | 1 | 1 (1) |  |  |  |
| Lepidoptera | Lycaenidae | 3 | 3 (2) |  | 1 (0) |  |
| Lepidoptera | Noctuidae | 74 | 75 (39) | 20 (1) | 25 (2) | 10 (0) |
| Lepidoptera | Nymphalidae | 10 | 10 (6) | 2 (0) | 4 (0) |  |
| Lepidoptera | Pieridae | 3 | 3 (3) |  |  |  |
| Lepidoptera | Pyralidae | 1 | 1 (1) |  |  |  |
| Lepidoptera | Sphingidae | 1 | 1 (1) |  |  |  |
| Lepidoptera | Tortricidae | 1 | 2 (1) |  |  |  |
| Lepidoptera | Zygaenidae | 4 | 3 (2) | 1 (0) | 1 (1) |  |
| Neuroptera | Chrysopidae | 10 | 7 (3) | 1 (0) | 4 (0) | 5 (3) |
| Neuroptera | Coniopterygidae | 3 | 3 (0) | 3 (1) | 1 (0) |  |
| Neuroptera | Hemerobiidae | 9 | 10 (4) | 5 (0) | 2 (0) | 2 (0) |
| Neuroptera | Sisyridae | 1 | 1 (1) |  |  |  |
| Opiliones | Nemastomatidae | 1 | 1 (0) | 1 (0) | 1 (0) | 1 (0) |
| Opiliones | Phalangiidae | 16 | 15 (2) | 10 (0) | 8 (0) | 13 (0) |
| Opiliones | Trogulidae | 1 | 1 (1) |  |  |  |
| Orthoptera | Acrididae | 9 | 9 (2) | 5 (0) | 7 (1) |  |
| Orthoptera | Phaneropteridae | 1 | 1 (0) |  | 1 (0) |  |
| Orthoptera | Tetrigidae | 2 | 2 (2) |  |  |  |
| Orthoptera | Tettigoniidae | 8 | 8 (1) | 6 (0) | 2 (0) | 3 (0) |
| Plecoptera | Taeniopterygidae | 1 | 1 (1) |  |  |  |
| Prostigmata | Eriophyidae | 4 | 4 (4) |  |  |  |
| Prostigmata | Erythraeidae | 1 |  | 2 (1) |  |  |
| Psocoptera | Amphipsocidae | 1 | 1 (0) | 1 (0) | 1 (0) |  |
| Psocoptera | Caeciliusidae | 8 | 9 (1) | 5 (0) | 4 (0) | 4 (0) |
| Psocoptera | Elipsocidae | 5 | 6 (2) | 4 (0) | 4 (0) | 2 (0) |
| Psocoptera | Lachesillidae | 1 | 1 (0) | 1 (0) | 1 (0) | 1 (0) |
| Psocoptera | Mesopsocidae | 3 | 3 (1) | 1 (0) | 2 (0) |  |
| Psocoptera | Peripsocidae | 5 | 6 (1) | 2 (0) | 4 (0) | 2 (0) |
| Psocoptera | Philotarsidae | 2 | 2 (0) | 1 (0) | 1 (0) | 2 (0) |
| Psocoptera | Psocidae | 8 | 7 (1) | 4 (0) | 6 (0) | 4 (1) |
| Psocoptera | Stenopsocidae | 2 | 3 (1) | 2 (0) | 2 (0) | 2 (0) |
| Psocoptera | Trogiidae | 2 | 2 (1) | 1 (0) | 1 (0) | 1 (0) |
| Raphidioptera | Raphidiidae | 2 | 2 (2) |  |  |  |
| Strepsiptera | Elenchidae | 1 | 1 (0) |  | 1 (0) | 1 (0) |
| Strepsiptera | Halictophagidae | 1 | 1 (1) |  |  |  |
| Trichoptera | Beraeidae | 3 | 2 (1) | 1 (1) |  | 1 (0) |
| Trichoptera | Ecnomidae | 1 | 1 (0) |  | 1 (0) |  |
| Trichoptera | Hydropsychidae | 2 | 1 (1) |  | 1 (1) |  |
| Trichoptera | Hydroptilidae | 7 | 7 (4) |  | 3 (0) |  |
| Trichoptera | Lepidostomatidae | 1 | 1 (1) |  |  |  |
| Trichoptera | Leptoceridae | 12 | 11 (8) | 2 (0) | 3 (1) |  |
| Trichoptera | Limnephilidae | 36 | 36 (15) | 15 (0) | 13 (0) | 11 (1) |
| Trichoptera | Molannidae | 2 | 2 (2) |  |  |  |
| Trichoptera | Phryganeidae | 6 | 6 (1) | 3 (0) | 2 (0) | 1 (0) |
| Trichoptera | Polycentropodidae | 6 | 6 (3) | 2 (0) | 1 (0) | 2 (0) |
| Trichoptera | Psychomyiidae | 3 | 3 (3) |  |  |  |
| Trichoptera | Sericostomatidae | 1 | 1 (0) |  |  | 1 (0) |

**Appendix G:** Species accumulation curves for arable sites (n=15), plantations (n=15) and natural sites (n=90).

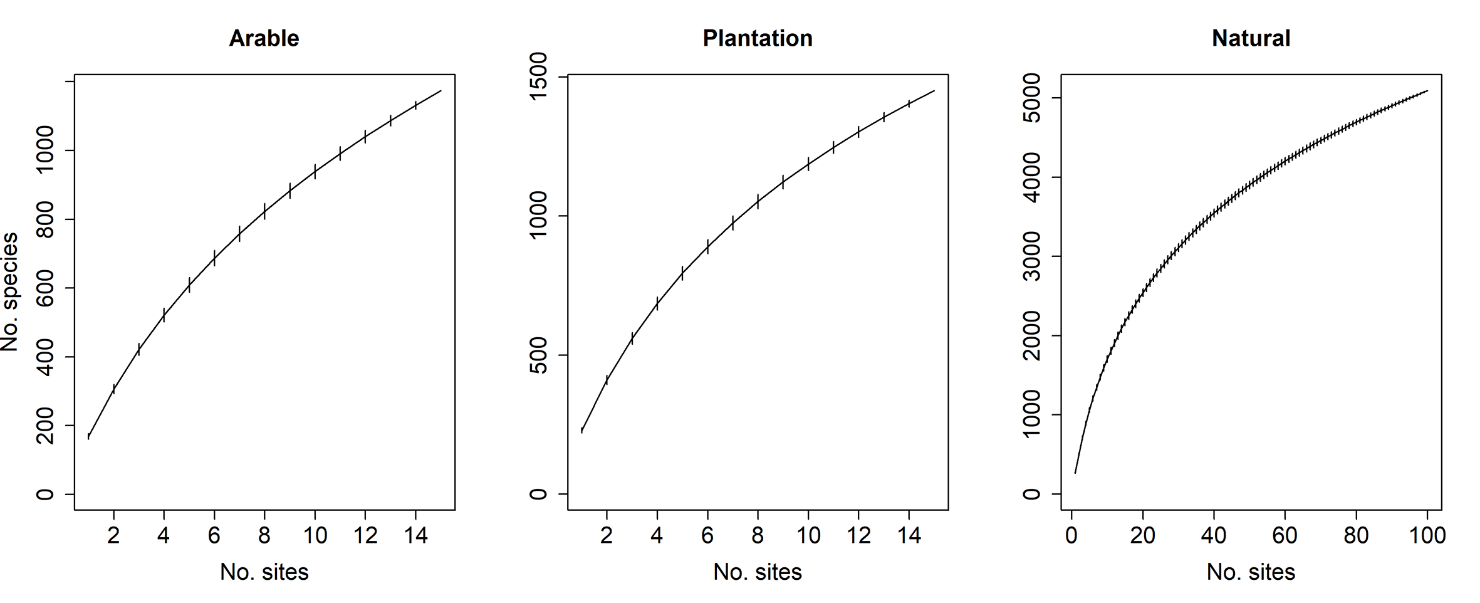

**Appendix H:** Spearman correlation (r) between measured abiotic and biotic environmental variables and Nonmetric Multidimensional Scaling (NMDS) plant and macrofungi axes 1-3. N = 130 for plant NMDS and N = 124 for macrofungi NMDS (sites with less than five species were omitted). Bold indicates significance at 0.05 level.

|  | PlantNMS1 | PlantNMS2 | PlantNMS3 | FungiNMS1 | FungiNMS2 | FungiNMS3 |
| --- | --- | --- | --- | --- | --- | --- |
| Litter mass | **-0.302** | **-0.730** | 0.040 | **0.652** | **0.341** | 0.106 |
| Soil pH | **0.661** | 0.079 | 0.026 | **-0.432** | **0.475** | 0.076 |
| Soil C | 0.058 | **0.189** | 0.028 | **-0.203** | -0.030 | 0.029 |
| Soil N | **0.580** | -0.002 | **0.255** | **-0.363** | **0.274** | **0.296** |
| Soil P | **0.733** | -0.103 | **0.205** | **-0.428** | **0.349** | **0.317** |
| Leaf N | **0.361** | **-0.529** | **-0.266** | 0.054 | **0.542** | -0.065 |
| Leaf C | **-0.507** | **0.368** | -0.066 | 0.063 | **-0.527** | **-0.198** |
| Leaf P | **0.509** | **-0.219** | -0.023 | **-0.246** | **0.376** | 0.175 |
| Deadwood volume | -0.141 | **-0.440** | 0.123 | **0.308** | **0.239** | 0.085 |
| Mean vegetation height | **0.240** | 0.036 | **-0.425** | -0.156 | **0.161** | **-0.321** |
| Trimmed mean soil moisture | **-0.211** | 0.121 | **-0.842** | -0.035 | **0.029** | **-0.703** |
| Max surface temperature | 0.063 | **0.583** | **0.228** | **-0.417** | **-0.376** | 0.147 |
| Sd air temperature | 0.105 | **0.495** | 0.139 | **-0.516** | **-0.267** | 0.102 |
| 85 quantile Vapour Pressure Deficit | 0.087 | **0.441** | **0.326** | **-0.392** | **-0.303** | **0.238** |
| Median light intensity | 0.026 | **0.723** | 0.071 | **-0.462** | **-0.493** | -0.171 |
| Plant species richness | **0.483** | **0.209** | -0.036 | **-0.430** | **0.358** | 0.032 |
| Trees > 40DBH (count) | -0.095 | **-0.672** | 0.059 | **0.479** | **0.343** | 0.092 |
| Trees > 40DBH (no. of species) | -0.149 | **-0.685** | 0.091 | **0.500** | **0.300** | 0.134 |
| Mean bare soil cover | -0.168 | **-0.695** | 0.139 | **0.617** | **0.336** | 0.088 |
| Mean bryophyte cover | **-0.386** | **0.303** | -0.039 | 0.034 | **-0.523** | -0.053 |
| Mean lichen cover | -0.026 | **0.206** | **0.313** | -0.109 | **-0.231** | 0.119 |
| Deadwood density | -0.034 | -0.147 | -0.146 | 0.099 | 0.133 | -0.092 |
| Anthill density | -0.076 | 0.124 | 0.073 | -0.018 | -0.128 | 0.063 |
| Boulder density | 0.082 | -0.075 | 0.170 | 0.023 | 0.065 | 0.129 |
| Water puddle density | 0.035 | -0.011 | **-0.206** | -0.010 | 0.095 | **-0.191** |
| Dung density | 0.056 | **0.269** | **0.234** | **-0.219** | -0.146 | **0.266** |
| Flower density | 0.082 | 0.149 | 0.155 | -0.122 | -0.016 | 0.173 |
| Temporal continuity class | **-0.209** | 0.007 | 0.095 | 0.100 | -0.103 | 0.010 |
| Spatial continuity 500m | **0.275** | -0.117 | **0.291** | 0.094 | -0.039 | **0.378** |
| Spatial continuity 1000m | **0.355** | -0.102 | **0.182** | 0.070 | -0.095 | **0.368** |
| Spatial continuity 2000m | **0.377** | -0.085 | 0.125 | 0.069 | -0.139 | **0.342** |
| Spatial continuity 5000m | **0.468** | -0.082 | 0.054 | -0.038 | -0.115 | **0.358** |
